# Robustness of high-throughput prediction of leaf ecophysiological traits using near infrared spectroscopy and poro-fluorometry

**DOI:** 10.1101/2025.02.07.636194

**Authors:** Eva Coindre, Romain Boulord, Laurine Chir, Virgilio Freitas, Maxime Ryckewaert, Thomas Laisné, Virginie Bouckenooghe, Maëlle Lis, Llorenç Cabrera-Bosquet, Agnès Doligez, Thierry Simonneau, Benoît Pallas, Aude Coupel-Ledru, Vincent Segura

## Abstract

Water scarcity is a major threat to crop production and quality. Improving drought tolerance through variety selection requires a deeper understanding of plant ecophysiological responses, but large-scale phenotyping remains a bottleneck. This study assessed the potential of high-throughput tools (spectroscopy and poro-fluorometry) to predict leaf morphological and ecophysiological traits in a grapevine diversity panel grown in pots under well-watered outdoor conditions and under three contrasting soil water treatments in a greenhouse. We found a certain complementarity between measuring devices. Spectrometers could accurately predict leaf mass per area, water content, and water quantity (R² > 0.58), while the poro-fluorometer was efficient for predicting net CO₂ assimilation (R² > 0.72), regardless of the water treatment. The prediction of leaf mass per area using spectrometers appeared to be quite robust across both outdoor and greenhouse experiments, while the prediction of water use efficiency was dependent on the water treatment, with much better predictions under moderate (R² > 0.73) than severe water deficit. Calibrated models were then applied to the full diversity panel using only high-throughput measurements to estimate trait values and their broad-sense heritability. Leaf mass per area, also measured directly, showed similar heritability whether based on observed or predicted data. Heritability estimates for predicted traits reached up to 0.5. Overall, our findings support the use of spectroscopy and poro-fluorometry as reliable, non-destructive tools for high-throughput phenotyping, enabling genetic studies on drought-related traits in grapevine.

## INTRODUCTION

Over the last decades, climate change has become a main concern. Prediction expects significant temperature rises and changes in annual rainfall patterns (Calvin et al., 2023; Semia Cherif et al., 2020; Van Leeuwen et al., 2024). In order to maintain crops in this context, there is a need to explore new combinations of genetic and management choices. Identifying the best performing solutions further requires high throughput characterization of traits related to plant functioning in responses to the environment.

Gas exchange, leaf structure and specific leaf biochemical components are among the traits most impacted by abiotic constraints such as soil water availability. The regulation of gas exchange at the leaf level by stomatal closure/opening plays a key role in the whole plant response to water stress. Indeed, a soil water deficit leads to a drop in water potential all along the plant that could damage its water transfer capacity and result in organ or plant death (Simonneau et al., 2017). To prevent such damage, plants can quickly close their stomata, reducing transpiration (E) (Damour et al., 2010) and maintaining their leaf water content (WC_f_) (Bhattacharya, 2021). However, this water saving strategy also decreases net photosynthesis (A_n_) by decreasing CO_2_ uptake (Simonneau et al., 2017), and in turn biomass production. A trade-off between reducing E and maintaining A_n_ under water stress requires considering water use efficiency, *i.e.* the biomass produced per unit of water consumed. At the leaf level, it can be measured by the instantaneous water use efficiency (WUE_inst_), calculated as the ratio between A_n_ and E, and the intrinsic water use efficiency (WUE_intr_), calculated as the ratio between A_n_ and the stomatal conductance for water vapor (g_sw_). The latter provides an estimate less dependent on air vapor pressure deficit than the former (Coupel-Ledru, 2021).

In grapevine, the species studied in this work, genotypic variability has already been reported for A_n_, g_sw_, WUE_intr_, and leaf water potential (Ѱ) (Bota et al., 2001; Tomás et al., 2014; Prieto et al., 2010). However, this variability was observed for a limited number of cultivars and/or in a limited number of growing conditions, usually well-watered conditions. High-throughput assessment of these ecophysiological traits is limited by the time required for measurements using leaf gas analyzers or pressure chambers. Studying large populations with a wide range of genetic diversity under various conditions is crucial to identify the best performing cultivars. It can also enable the identification of genetic regions associated with these traits and their responses to contrasted environmental conditions (Coupel-Ledru et al., 2014; Trenti et al., 2021). A main challenge is thus to investigate the use of alternative high-throughput methods as proxies to accurately predict traits of interest on large panels of genotypes under various water availability treatments and under different growing conditions (Xiao et al., 2022).

Fluorometer devices, providing chlorophyll fluorescence parameters related to photosystem II efficiency and electron transport rate, have been proposed as low-cost and rapid tools to estimate An (Murchie & Lawson 2013; Han et al., 2022). Chlorophyll fluorescence measurements were also reported to evaluate the variability of photosynthetic performance among a population of hybrids in barley (Fernández-Calleja et al., 2020). Moreover, previous studies have calculated a semi-empirical index that combines fluorescence parameters and leaf temperature, to estimate net photosynthesis in fruit trees (Losciale et al., 2015). This indicator was then used to evaluate the photosynthesis activity of a core-collection of apples in response to water deficit (Coupel-Ledru et al., 2019). Near-infrared spectroscopy (NIRS) is another high-throughput method commonly used to predict plant tissue composition (Foley et al., 1998; Petisco et al., 2006; Cabrera-Bosquet et al., 2012). Dedicated chemometrics methods, such as Partial Least Square Regression (PLSR) (Wold et al., 2001), are used to establish relationships between spectral absorbance or reflectance and ground-truth data obtained by low-throughput methods. NIRS has been reported in plants to predict numerous leaf compositional traits, including nitrogen (Ge et al., 2019) and carbon contents, lignin, cellulose (Petisco et al., 2006), protein, starch, total non-structural carbohydrates (De Bei et al., 2017) and sugars concentrations (Ely et al., 2019; Van Wyngaard et al., 2021). NIRS has also been implemented to predict various morphological and ecophysiological traits on multiple species and on large populations of the same species. For instance, leaf mass per area (LMA), which is widely used in plant ecology as an indicator of plant functioning, has been predicted by NIRS on diverse species (Ely et al., 2019; Serbin et al., 2019). NIRS predictive models were also reported for LMA and A_n_ on wheat and triticale germplasms (Silva-Perez et al., 2018) as well as on hybrids of maize (Cotrozzi et al., 2020). LMA, A_n_, WUE_intr_, WUE_inst_, Ѱ and WC_f_ were together predicted by NIRS on a diversity panel of maize (Ge et al., 2019) and Ѱ has also been predicted by NIRS on several cultivars of grapevine (De Bei et al., 2011; Rapaport et al., 2015). The relevance of NIRS to predict water related traits was observed to be associated with light absorption by water at wavelength bands centered around 1400 and 1900 nm (Workman Jr. & Weyer, 2012; Das et al., 2021). To minimize the impact of water on spectra and consequently on the prediction accuracy of other traits, spectra can be collected on dried leaves (Masemola & Cho, 2019). Finally, not all spectrometers measure the same wavelength range. Therefore, the type of spectrometer must be rationalized according to the application (Zhu et al., 2022; Barthès et al., 2019).

When studying ecophysiological traits, the robustness of PLSR model predictions must be examined. On grapevine, predictive models with NIRS data have been built on few cultivars subjected to different soil water treatments (De Bei et al., 2011; Rapaport et al., 2015; Ryckewaert et al., 2022a). PLSR models were calibrated and validated with datasets containing samples from all water treatments (De Bei et al., 2011; Ryckewaert et al., 2022a), without assessing the ability of the model calibrated on a given water treatment to predict the values of plants subjected to another soil water treatment. If PLSR models are robust across different aerial environmental conditions and soil water availabilities, phenotyping time could be reduced in new experiments. Another issue with the PLSR model is the risk of overfitting. To prevent this risk, the calibration model is usually built with a cross-validation procedure. However, even in a cross-validation setting, the predictive ability of the model could still depend on the calibration and validation sets that could be randomly or non-randomly chosen (Filzmoser et al., 2009).

In this study, we evaluated the ability and complementarity of NIRS and poro-fluorometry data to predict leaf traits at high-throughput. These two approaches were applied on a grapevine diversity panel (Nicolas et al., 2016), grown in pots both outdoors under non-limiting water availability and in a greenhouse with varying intensities of soil water deficit. The robustness of PLSR models was studied across experiments and water treatments. Heritabilities of traits either directly measured or estimated from high-throughput data were compared to investigate the ability of high-throughput phenotyping to assess the genotypic variability.

## MATERIALS AND METHODS

### Plant material and experimental design

We used a subset of 241 cultivars from a panel of 279, previously designed to represent the worldwide genetic diversity of *Vitis vinifera* (Nicolas et al., 2016). We added 5 cultivars that are in the process of being registered in the French catalog, resulting in 246 studied cultivars. Twenty own-rooted plants were grown for each cultivar with ferti-irrigation from 2018 at the experimental vineyard of l’Institut Agro Montpellier (France) in individual 3-liter pots containing a 30:70 (v/v) mixture of loamy soil and organic compost (Coupel-Ledru et al., 2024). In January 2020, the plants were transferred into larger, individual 9-liter pots filled with the same substrate.

A first phenotyping season was conducted in July 2021, on the potted plants grown in the experimental vineyard under non-limiting, drip-irrigated conditions (“Outdoor experiment”). Six plants out of the 20 available per cultivar were next retained to be part of a greenhouse experiment in 2022, where different watering treatments were set up (“Greenhouse experiment”). This second experiment was conducted in the PhenoArch phenotyping platform (Cabrera-Bosquet et al., 2016) hosted at theMontpellier Plant Phenotyping Platforms (https://eng-lepse.montpellier.hub.inrae.fr/platforms-m3p/montpellier-plant-phenotyping-platforms-m3p), from May to August 2022. The whole panel (246 cultivars) was subjected to three different water treatments: Well-Watered (WW), moderate Water Deficit (WD1) and severe Water Deficit (WD2), with two plants for each cultivar and each water treatment (1476 plants in total). These treatments corresponded to soil water potential equal to -0.03 MPa (well watered conditions, WW), -0.11 Mpa (moderate water deficit, WD1) and -0.43 Mpa (severe water deficit, WD2) (protocol for management of water treatment as in Coupel-Ledru et al., (2014)). For each cultivar, the six plants were grouped into two triplets (one triplet including one WW, one WD1 and one WD2 plant), and cultivar triplets were randomized within the phenotyping platform. Environmental variables were monitored using 10 probes, distributed throughout the greenhouse.

### Leaf phenotyping

Leaf phenotyping was performed on two to three plants per cultivar in the outdoor experiment while all the individuals were phenotyped in the greenhouse experiment once the water content in the pots had stabilized at their target values. For both experiments (outdoors and greenhouse), measurements were performed between 9:00 and 13:00 (UTC+2). Leaves were chosen as fully expanded and well exposed to sunlight. They were usually located between the tenth and the fifteenth phytomer from the apex. Two kinds of measurements were performed on selected leaves: high- and low-throughput measurements. High-throughput measurements were carried out during 5 mornings on a total of 838 leaves in the outdoor experiment in July 2021 and during 24 mornings on a total of 1450 leaves in the greenhouse experiment in June and July 2022. Low-throughput measurements were performed only in the greenhouse experiment on a subset of 120 plants selected to be representative of the experimental design and thus covering the three watering conditions.

Multiple devices were used successively on the same leaf *in situ*, in a short time interval, for both high- and low-throughput phenotyping. The first device used was a hand-held poro-fluorometer (Li600, Portable Photosynthesis System LI-600, Licor, Lincoln, United States). It was used to measure the transpiration rate, E (mmol m^−2^ s^−1^), the stomatal conductance g_sw_ (mol m^−2^ s^−1^), chlorophyll fluorescence parameters and micro-climate variables i.e. leaf vapor pressure deficit, atmospheric pressure and ambient light (Supplemental Table S1). Two measurements were taken at two different points on the same leaf and were averaged to obtain a single value per leaf. The second and third devices were micro-portable near infrared spectrometers: MicroNIR (MicroNIR™ Onsite-W, VIAVI, Chandler, AZ, United States) and NeoSpectra (NeoSpectra™ Scanner, Si-Ware, Miami, FL, United States). The MicroNIR covers a spectral range from 908 nm to 1676 nm with 125 wavelengths, resulting in a spectral sampling interval of 6.2 nm. The NeoSpectra covers a spectral range from 1350 nm to 2500 nm with 257 wavelengths and a spectral sampling interval of 16 nm. The wavelength interval between 1350 nm and 1676 nm was common between both devices. For both devices, two spectra were collected per leaf on the lamina while avoiding the principal veins. These two spectra were respectively collected on each side of the leaf (abaxial and adaxial faces) and were then averaged to obtain a single spectrum per leaf. After these measurements, a 3 cm diameter disk was sampled from the leaf. In the greenhouse experiment, this disk was immediately weighed after sampling to obtain its fresh weight (FW, mg). For both outdoor and greenhouse experiments, the disk was oven-dried for one week at 60°C before being weighed to get its dry weight (DW, mg). Leaf Mass per Area (LMA, mg cm^−^²) was calculated as the ratio between DW and disk area. For disks of leaves collected in the greenhouse experiment only, leaf water content (WC_f_, mg H_2_O mg^−1^ fresh weight) was also calculated, as (FW - DW) / FW and the water quantity (WQ, mg H_2_O per leaf disk) as (FW-DW). After weighing, dried disks were stored at 20°C. They were later used again to collect NIRS spectra using a laboratory spectrometer ASD (ASD LabSpec® 4, Analytical Spectral Devices, Boulder, USA). The temperature was maintained at 20°C in the lab during measurements. Four spectra were taken using a reflectance probe (4 mm diameter) at different points of the dried disk before being averaged. The ASD covers a spectral range from 350 nm to 2500 nm with 2150 wavelengths and a sampling interval of 1 nm.

The low-throughput measurements were performed only in the greenhouse experiment during 4 mornings. The measurements started as described above for the high-throughput devices (i.e. Li600, MicroNir, NeoSpectra), and were followed by a measure with an InfraRed Gas-exchange Analyzer (IRGA) equipped with a light-emitting diode light source (Li6800, Portable Photosynthesis System, Licor, Lincoln, United States). This device allowed measuring net CO_2_ assimilation (A_n_, µmol m^−2^ s^−1^), transpiration rate (E, mmol m^−2^ s^−1^) and stomatal conductance (g_sw_, mol m^−2^ s^−1^). Intrinsic Water Use Efficiency (WUE_intr_, µmol CO_2_ mol^−1^ H_2_O) and instantaneous Water Use Efficiency (WUE_inst_, µmol CO_2_ mmol^−1^ H_2_O) were further calculated as the ratio between A_n_ and g_sw_ or A_n_ and E, respectively. The conditions in the leaf chamber during the measurement were set to 400 µmol mol^−1^ CO_2_, with actinic light adjusted to a value corresponding to the ambient Photosynthetically Photon Flux Density (PPFD) measured simultaneously by the Li600 device for each measured leaf. PPFD values thus ranged from 110 to 1150 µmol m^−2^ s^−1^ in the entire dataset. The leaf water potential (Ѱ, MPa) was next measured on these leafs with a Scholander pressure (SoilMoisture Equipment Corp., Goleta, United States).

### Preprocessing of spectral data

All data analyses were handled in R v4.3.0 (R Core Team, 2023). Spectra from the ASD laboratory spectrometer were initially processed using the rmgap function (R/rchemo package) (Brandolini-Bunlon et al., 2023) to remove vertical gaps caused by differences between detectors. Wavelengths between 350 and 400 nm were removed and excluded from spectral analyses due to their noisy signals (Ecarnot et al., 2013). The spectra were then smoothed using R/loess, with the neighborhood size controlled by an α value of 0.03. In order to express spectra from all devices in reflectance and wavelength (nm), the MicroNIR absorbance (A) was converted into reflectance (R), as R=1/10^A^, and the NeoSpectra wavenumber (ν, cm^−1^) was converted into wavelength (λ, nm), as λ=10^7^/ν. As the MicroNIR and NeoSpectra spectrometers have complementary ranges, with an overlap from 1350 nm to 1676 nm, a new, additional spectrum called MicroNIR_NeoSpectra was created for each leaf. Whole MicroNIR spectra from 950 nm to 1676 nm were retained due to their higher resolution than NeoSpectra spectra and combined with the 1676 nm - 2500 nm ranges measured with the NeoSpectra. The resulting gap at the junction of the two spectra was removed using R/rmgap from R/rchemo. All spectra (ASD, MicroNIR, NeoSpectra and MicroNIR_NeoSpectra) were then preprocessed using Standard Normal Variate (SNV) (Barnes et al., 1989). This was followed by a first-order derivative using the Savitzky-Golay filter (R/sgolayfilt from R/signal package (signal developers, 2023; Savitzky & Golay, 1964)). The polynomial order for the first-order derivative was set to 2, and the window sizes were set to 11, 15, 15 and 41 for MicroNIR, NeoSpectra, MicroNIR_NeoSpectra and ASD, respectively, to account for their differences in resolution.

### Data modeling and cross-validation

Calibration models were constructed in order to relate high-throughput data from the spectrometers on the one hand and twenty-one variables measured with the poro-fluorometer Li600 (Supplemental Table S1) on the other hand to the traits of interest measured with low-throughput devices or methods (IRGA, pressure chamber, disks weighing) using PLSR, as implemented in R/pls with the “kernelpls” method (Liland et al., 1999). The considered traits of interest were LMA, WC_f_, WQ, A_n_, Ѱ, WUE_intr_ and WUE_inst_, using data from the greenhouse experiment. For the outdoor experiment, only LMA was considered as the other traits were not measured.

A total of 11 datasets were designed to address specific research questions, while accounting for the experimental design and available data (Table 1). The first 2 datasets (A and B) aimed at evaluating the potential and complementarity of high-throughput phenotyping tools (spectrometers and poro-fluorometer Li600) for predicting traits of interest. These datasets included the maximum number of available data from the greenhouse experiment. Model training and evaluation were jointly carried out through a 5-fold cross-validation repeated 10 times and based on cultivars rather than individual leaves to avoid the confounding effect of cultivar prediction on trait prediction. The next 3 datasets (C, D and E) were designed to assess model robustness across greenhouse and outdoor experiments for the prediction of LMA (the only trait available in both experiments), while the following 4 datasets (F, G, H and I) were designed to assess model robustness across water treatments (within the greenhouse experiment) for the prediction of LMA, WC_f_ and WQ. All these datasets (C to I) included a fixed number of 370 leaves for the calibration, which were evenly and randomly sampled within the different experimental conditions: outdoors (C), greenhouse (D), outdoors and greenhouse (E), WW (F), WD1 (G), WD2 (H), the 3 water treatments (I). Model training for these calibration sets was done through 4-fold cross-validations repeated 10 times. Model evaluations were carried out within independent validation sets composed of random samples of 100 leaves repeated 15 times and specifically designed to evaluate prediction accuracy with respect to the experiment (greenhouse, outdoors) or the water treatment (WW, WD1, WD2). Dataset J was designed to evaluate model robustness across water treatments within the greenhouse experiment for the prediction of A_n_, Ѱ, WUE_intr_, WUE_inst_. This dataset was set to include a total of 72 and 30 leaves in the calibration and validation sets, respectively. These leaves were equally distributed in each water treatment. Since the number of leaves within the calibration set was rather low, model calibration for this dataset was done with leave-one-out (LOO) cross-validation. Finally, a dataset (K) was designed for predicting traits of interest within the greenhouse experiment and further assess the genetic variability of predicted values (see hereafter). This dataset included a total of 102 leaves within the calibration set evenly distributed per water treatment, and in this case model training was done through a 4-fold cross-validation scheme.

**Table 1.** Strategies to evaluate the robustness and the prediction quality of PLSR models.

| Rationale | Dataset | LT or HT | Traits | Calibration |  |  |  |  |  |  | Validation |  |  |  |  |
| --- | --- | --- | --- | --- | --- | --- | --- | --- | --- | --- | --- | --- | --- | --- | --- |
|  |  |  |  | Number of leaves |  |  |  |  | Cross-validation settings |  | Number of leaves |  |  |  | Settings |
|  |  |  |  | WW | WD1 | WD2 | Outdoors | Total | Nb fold | Nb iter. | WW | WD1 | WD2 | Outdoors | Nb iter. |
| Devices complementarity | **A** | HT | LMA, WC<sub>f</sub>, WQ | 483 | 481 | 473 | - | 1437 | 5-fold | 10 | - | - | - | - | - |
| | **B** | LT | LMA, A<sub>n</sub>, $\Psi$, WUE<sub>intr</sub>, WUE<sub>inst</sub> | 37 | 36 | 36 | - | 109 | 5-fold | 10 | - | - | - | - | - |
| Robustness between experiments | **C** | HT | LMA | - | - | - | 370 | 370 | 4-fold | 10 | 100 | 100 | 100 | 100 | 15 |
|  | **D** | HT | LMA | 124 | 123 | 123 | - | 370 | 4-fold | 10 | 100 | 100 | 100 | 100 | 15 |
|  | **E** | HT | LMA | 93 | 92 | 93 | 92 | 370 | 4-fold | 10 | 100 | 100 | 100 | 100 | 15 |
| Robustness between water treatments | **F** | HT | LMA, WC<sub>f</sub>, WQ | 370 | - | - | - | 370 | 4-fold | 10 | 100 | 100 | 100 | - | 15 |
|  | **G** | HT | LMA, WC<sub>f</sub>, WQ | - | 370 | - | - | 370 | 4-fold | 10 | 100 | 100 | 100 | - | 15 |
|  | **H** | HT | LMA, WC<sub>f</sub>, WQ | - | - | 370 | - | 370 | 4-fold | 10 | 100 | 100 | 100 | - | 15 |
|  | **I** | HT | LMA, WC<sub>f</sub>, WQ | 124 | 123 | 123 | - | 370 | 4-fold | 10 | 100 | 100 | 100 | - | 15 |
| | **J** | LT | A<sub>n</sub>, $\Psi$, WUE<sub>intr</sub>, WUE<sub>inst</sub> | 24 | 24 | 24 | - | 72 | LOO | 10 | 10 | 10 | 10 | - | 100 |
| Predictive models | **K** | LT | LMA, A<sub>n</sub>, WUE<sub>intr</sub>, WUE<sub>inst</sub> | 34 | 34 | 34 | - | 102 | 4-fold | 1 | - | - | - | - | 1 |

Leaf mass per area (LMA), Water content (WC<sub>f</sub>), Water quantity (WQ), Net CO<sub>2</sub> assimilation (A<sub>n</sub>), Leaf water potential ( $\Psi$ ), Intrinsic water use efficiency (WUE<sub>intr</sub>), Instantaneous water use efficiency (WUE<sub>inst</sub>), High-throughput phenotyping (HT), Low-throughput phenotyping (LT), Number of iterations (Nb Iter.), Leave-one out (LOO), Well-watered treatment (WW), Moderate water deficit treatment (WD1), Severe water deficit treatment (WD2).

Whatever the dataset, PLSR models were always built with a maximum number of latent variables set at 20 for spectrometers and 10 for the Li600. The optimal number of latent variables was determined by minimizing the average Root Mean Squared Error (RMSE) over the iterations of the repeated cross-validations (Datasets A to J) or directly the RMSE for the LOO cross-validation (Dataset J). Model quality was evaluated either within the cross-validation (Datasets A and B) or using validation (testing) sets (Datasets C to J) using the R^2^ computed as the squared correlation between observed and predicted values, the RMSE or its normalized version to allow comparisons between traits (NRMSE). Significant differences between model performances across experimental conditions or water treatments were assessed with a Tukey HSD post-hoc test, as implemented in R/stats.

### Heritability of predicted traits

PLSR models (R²_cv_ ≥ 0.55) calibrated with the dataset K were used to predict traits within the greenhouse experiment using high-throughput measurements (fluo-porometers and NIRS data). The following mixed-linear model was fitted to the predicted values per water treatment in order to decompose their phenotypic variation using R/lme4 (Bates et al., 2015):

Y_ij_ = µ + G_i_ + ΣA_ij_ + ɛ_ij_, where Y_ij_ represented the vector of predicted values per water treatment, µ was the intercept, G_i_ was the random effect of the cultivar i, ΣA_ij_ was a sum of fixed effects of experimental covariates measured on the plant j from the cultivar i and ɛ_ij_ was the random residual effect. The fixed effects included date of measurement, time of measurement, time elapsed between the last irrigation of the plant and the measurement, duration between the entrance of the plant into the platform and the measurement, environmental conditions as the average of the 10 probes (temperature, PPFD, relative humidity, vapor pressure deficit), and plant density around each plant. For each trait within each water treatment a model selection was carried to retain only the significant fixed effects (α = 0.05) using R/rutilstimflutre (Flutre & Chabrault, 2019). Variance components in the selected model were used to estimate broad-sense heritability per water treatment as: H²= σ²_G_ / (σ²_G_ + σ²_E_/n) with σ²_G_ the genotypic variance, σ²_E_ the residual variance and n the mean number of repetitions (leaves) per cultivar within each water treatment, typically around 2.

## RESULTS

### Variability of leaf traits

The lowest coefficient of variation was observed for WC_f_ (<5%), regardless of the water treatment. LMA and WQ showed intermediate coefficients of variation (between 15% and 22%), with generally slight differences between water treatments and experiments (Supplemental Table S2). LMA and WQ were positively correlated, with their correlation coefficient increasing from 0.63 to 0.73 with the intensity of water deficit in the greenhouse experiment (Supplemental Figure S1a, Supplemental Figure S1b, Supplemental Figure S1c). LMA and WC_f_ were negatively correlated, regardless of the water treatment, with correlation coefficient decreasing from -0.57 to -0.48 when water deficit increased. The correlation between WC_f_ and WQ was lower but significantly positive, with values close to 0.25 (Supplemental Figure S1a, Supplemental Figure S1b, Supplemental Figure S1c).

A_n_,Ѱ, WUE_intr_ and WUE_inst_ displayed more variability than LMA, WC_f_ and WQ (Supplemental Figure S2), as underlined by higher coefficients of variation (Supplemental Table S2). A_n_ was the trait whose coefficient of variation was the most affected by the water treatment, increasing with the intensity of water deficit. Conversely, the coefficient of variation of WUE_intr_ decreased with increasing water deficit. A_n_ was not significantly correlated with WUE_intr_ and WUE_inst_, whatever the water treatment (Supplemental Figure S1d, Supplemental Figure S1e, Supplemental Figure S1f). WUE_intr_ was strongly correlated with WUE_inst_ with values around 0.8 in all conditions (Supplemental Figure S1d, Supplemental Figure S1e, Supplemental Figure S1f).

### Devices complementarity for the prediction of leaf traits

Overall, spectra of the three spectrometers collected in the greenhouse experiment (Datasets A and B, Table 1) were highly effective in predicting traits such as LMA, WC_f_ and WQ. The coefficient of determination computed within the cross-validation procedure consistently exceeded 0.67 (Fig 1a, Supplemental Figure S3a, Supplemental Figure S3b, Supplemental Figure S3c) with almost no variability across the cross-validation repetitions. Nevertheless, significant differences in R²_cv_ were found between devices (Figure 1, Supplemental Table S3). The ASD spectrometer on dry leaves yielded the most accurate models to predict LMA compared to micro-portable spectrometers used *in situ* (Figure 1, Supplemental Figure S4a). Conversely, the best models for predicting WC_f_ and WQ were established with spectra from micro-portable spectrometers collected *in situ* (MicroNIR, NeoSpectra and MicroNIR_NeoSpectra in Figure 1a). Although model performances were almost similar for these three modalities, slightly better predictions were achieved by combining the wavelengths from MicroNIR and NeoSpectra. The PLSR models built with the variables from the Li600 device were unable to predict LMA, WC_f_ and WQ (R²_cv_ ≤ 0.1).

**Figure 1.**
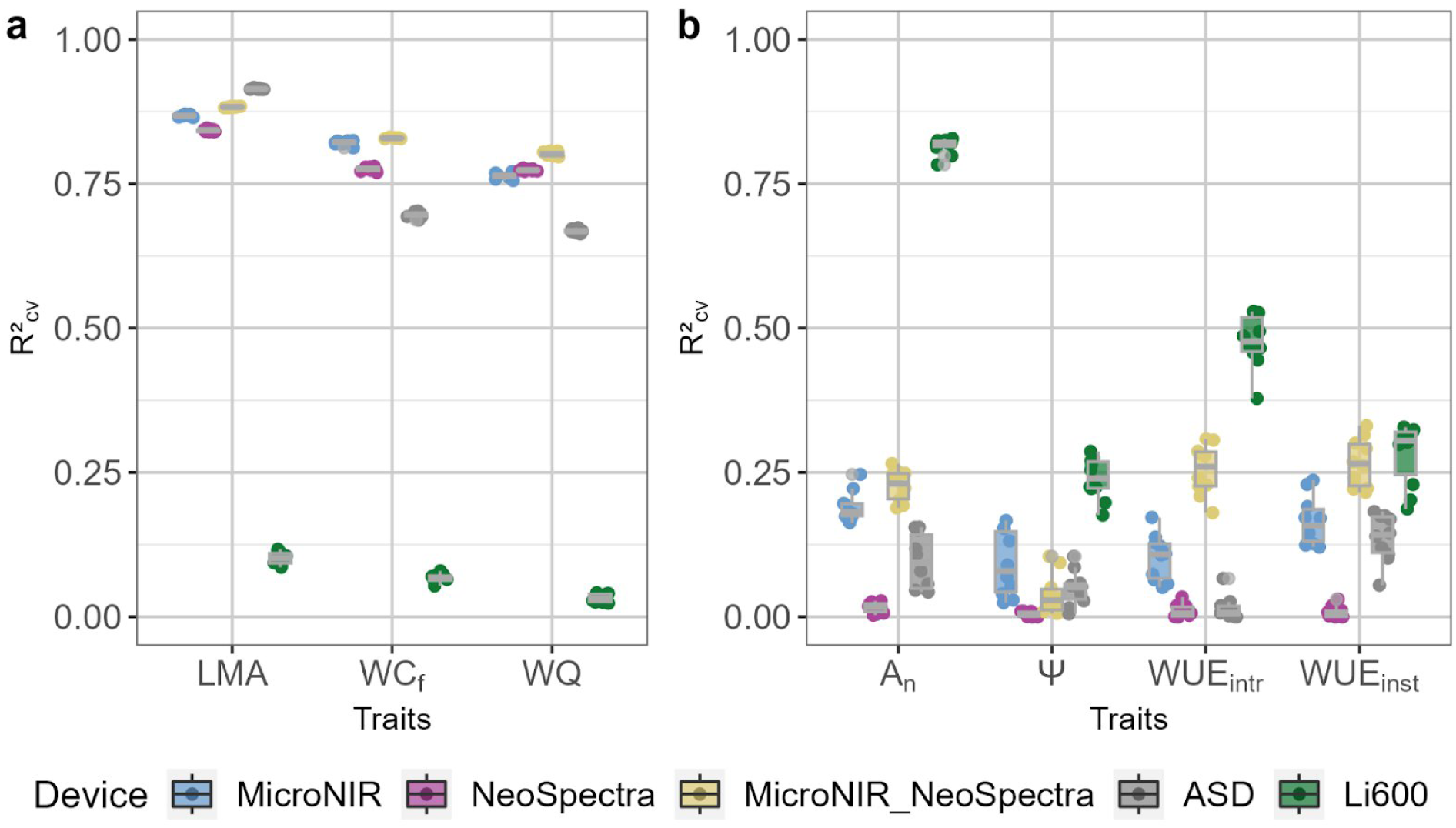
R²cv values computed in the cross validation procedure of leaf traits according to the device. a. High-throughput measured traits: leaf mass per area (LMA), Water content (WC_f_) and Water quantity (WQ) (Dataset A in Table 1). b. Low-throughput measured traits: Net CO2 assimilation (A_n_), Leaf water potential (Ѱ), Intrinsic water use efficiency (WUEintr) and Instantaneous water use efficiency (WUEinst) (Dataset B in Table 1). Each point in the boxplot corresponds to the R²_cv_ of one of the 10 iterations of the cross-validation (as detailed in Table 1).

Regarding the prediction of A_n_, Ѱ, WUE_intr_ and WUE_inst_ with spectrometers, R² values remained low and not higher than 0.27 whatever the device considered (Figure 1b). In contrast, the models built with variables from the Li600 device for predicting A_n_ displayed an R²_cv_ of 0.82 (Figure 1b, Supplemental Figure S3d, Supplemental Figure S4b). The Li600 device also seemed to capture relevant information for predicting WUE_intr_ and WUE_inst_, with R²_cv_ values of 0.48 and 0.3, respectively (Figure 1b, Supplemental Figure S3f, Supplemental Figure S3g). The prediction accuracies of WUE_inst_ using either the Li600 or the MicroNIR_NeoSpectra were not significantly different (Figure 1b, Supplemental Table S3). Ѱ was poorly predicted, regardless of the device used, with an R²_cv_ always below 0.24 (Figure 1b, Supplemental Figure S3e). The large variability in R²_cv_ or NRMSE_cv_ between the different cross-validation sets was likely due to the small number of leaves included in each set (dataset B in Table 1).

To further investigate the effect of calibration set size on model accuracy, we compared PLSR models built with low-(109 leaves) and high-throughput (1437 leaves) measurements of LMA. As expected, models built within the low-throughput calibration set displayed significantly lower R²_cv_ than models built within the high-throughput calibration set (by 0.08 to 0.15) (Supplemental Figure S5a). This might partly be due to a lower range of variation in the low-throughput calibration set compared to the high-throughput calibration set (Supplemental Figure S1h). This point is confirmed by the RMSE_cv_ values which displayed low differences between low- and high-throughput calibration sets with values always lower than 0.42 mg cm^−^² (Supplemental Figure S5b).

### Prediction robustness across experiments

The prediction accuracy for LMA was significantly lower within the outdoors than the greenhouse validation set (Figure 2), with median R² values ranging from 0.30 to 0.86 within the outdoors validation set and from 0.78 to 0.90 within the greenhouse validation set whatever the device considered. The ASD device used on dried disks yielded the most robust models for predicting LMA, regardless of the calibration and validation sets (Figure 2). Conversely the models built with micro-portable spectrometers on fresh leaves, especially the MicroNIR, appeared to be impacted by the environmental conditions in which they were calibrated and validated (Figure 2). This was particularly the case when models were calibrated with greenhouse data and validated on data collected outdoors. For MicroNIR, NeoSpectra and MicroNIR_NeoSpectra devices, the performance of models to predict LMA in outdoors conditions was similarly good whether calibrated with outdoor or greenhouse_outdoor calibration sets (Figure 2). Models calibrated with outdoors data remained accurate to predict greenhouse data since R² was greater than 0.78 and RMSE lower than 0.8 mg cm-². However, the prediction of greenhouse data, using an outdoors calibration set significantly reduced R² and RMSE values compared to greenhouse or greenhouse_outdoor calibration sets. This might partly be due to a lower coefficient of variation for LMA in outdoors (16.7) than in greenhouse (between 20.5 and 21.4) experiments (Supplemental Table S2, Supplemental Figure S2h, Supplemental Figure S2i).

**Figure 2.**
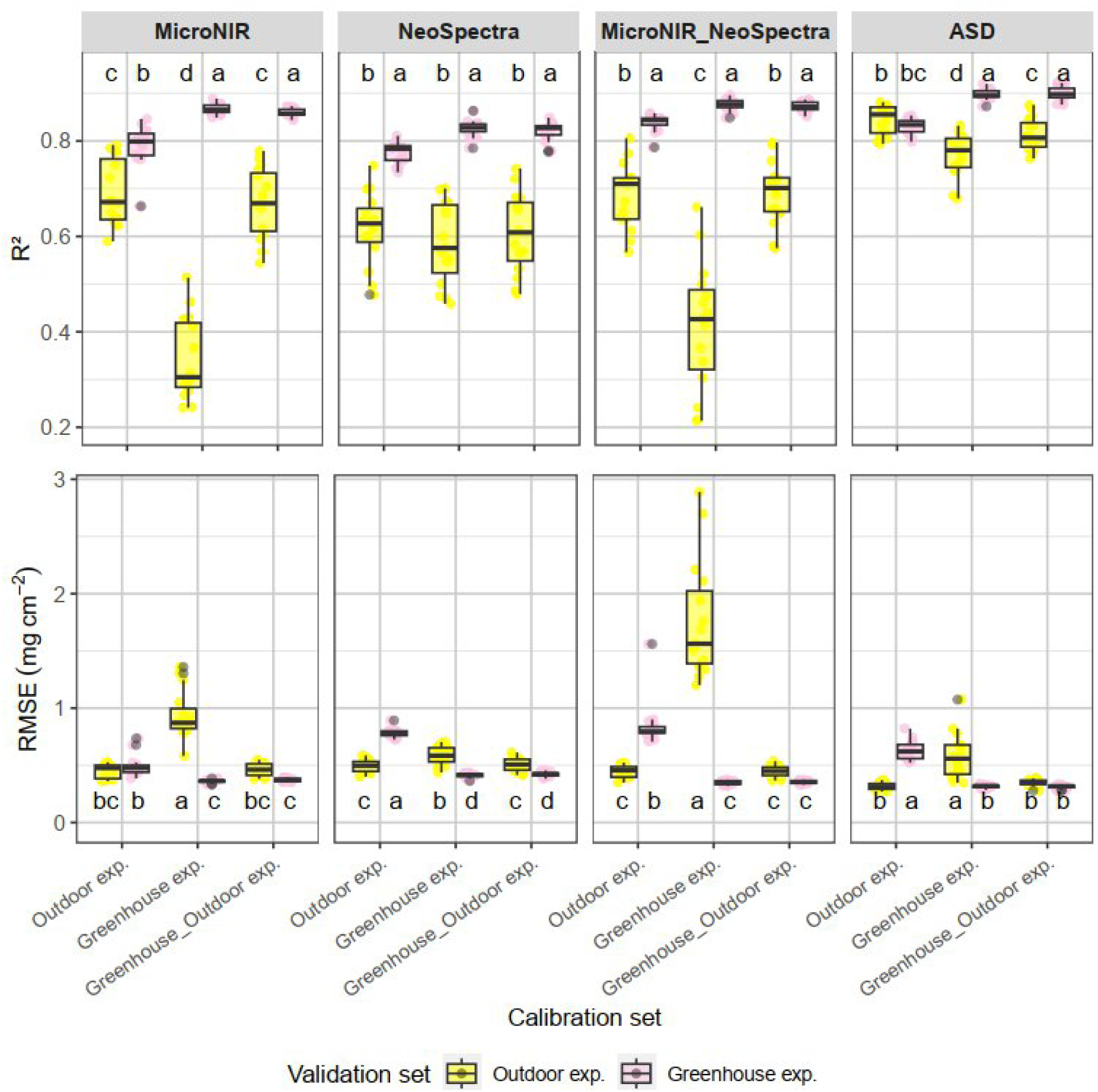
Prediction quality of LMA depending on whether spectra were collected in outdoors or greenhouse experiments. The top panel corresponds to the R² and the bottom panel corresponds to the RMSE (in mg cm-2) of the prediction. Each point in the boxplot represents one of the 15 iterations of the validation (Datasets C, D, E in Table 1). Different calibration sets are highlighted on the x-axis of each panel. Boxplots with the same letters within one facet are not significantly different (p > 0.05). exp.: experiment.

### Prediction robustness between soil water treatments

PLSR models were compared between water treatments for LMA, WC_f_ and WQ as these 3 traits were available on the entire diversity panel (Datasets F to J in Table 1). Partitioning the validation set according to the water treatment had more influence on model performance than partitioning the calibration set (Supplemental Figure S6). We thus decided to focus our analysis on the comparison of PLSR models calibrated with leaves from the 3 water treatments (Figure 3).

**Figure 3.**
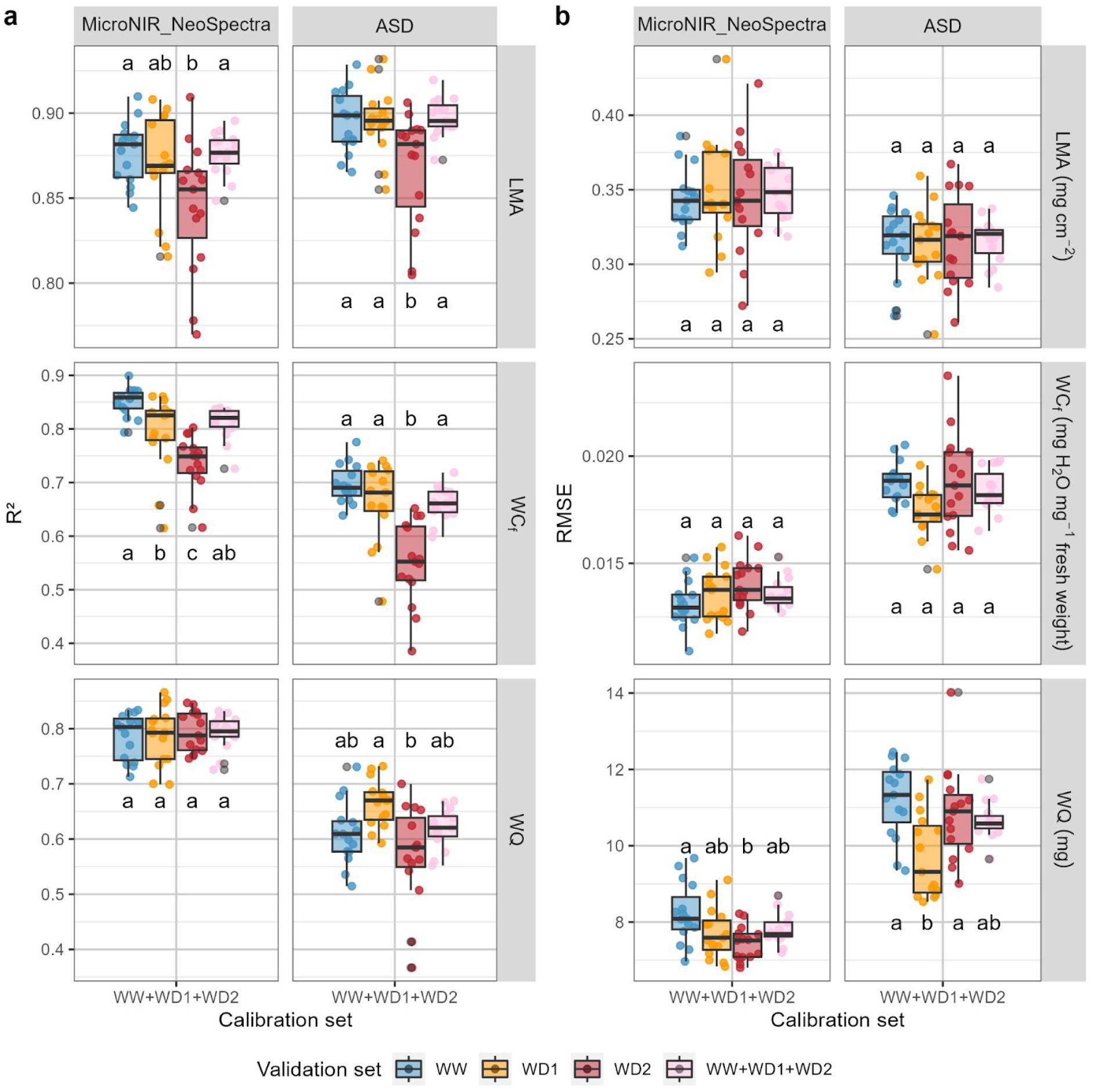
Prediction quality of PLSR models for LMA, WCf and WQ depending on the validation set. Models were calibrated using the WW+WD1+WD2 calibration set (Dataset I in Table 1) and the MicroNIR_NeoSpectra and ASD devices. a. corresponds to the R² and b. corresponds to the RMSE in the validation set. Each point in the boxplot represents one of the 15 iterations of the validation (as detailed in Table 1). The blue dots and boxes correspond to the well-watered treatment (WW); orange dots and boxes to the moderate water deficit treatment (WD1); red dots and boxes to the severe water deficit treatment (WD2); and pink dots and boxes to all WW, WD1 and WD2 treatments. LMA: leaf mass per area; WC_f_: water content; WQ: water quantity.

R² of predictive models was more impacted by the water treatment (Figure 3a) than RMSE (Figure 3b). For LMA, R² was significantly lower within WD2 compared to WW and WD1 validation sets (Figure 3a). These differences in R2 could be related to a decrease in the range of variation with the increase in water deficit (Supplemental Figure S2a), since the corresponding RMSE values did not significantly differ between water treatment (Figure 3b). Similarly for WC_f_ prediction with MicroNIR_NeoSpectra, R² significantly decreased from 0.86 to 0.75 when water deficit increased, while RMSE did not significantly vary (Figure 3a, Figure 3b). This influence of water treatment on trait prediction was not observed for WQ, with R² values always close to 0.8 when using the MicroNIR_NeoSpectra (Figure 3a) and median RMSE values lower than 8.1 mg H2O, regardless of the water treatment (Figure 3b). In contrast, the ASD device, used on dried leaves, yielded a less accurate prediction of WQ with lower R² values and RMSE always higher than 9.3 mg H2O. For this device prediction accuracy was significantly lower for the WD2 treatment as observed for LMA and WC_f_.

Because A_n_, Ѱ, WUE_intr_ and WUE_inst_ were phenotyped on a limited subset of leaves from the diversity panel, PLSR models were calibrated by gathering all water treatments, and they were evaluated separately for each water treatment (Dataset J in Table 1). The high variability of model performances observed with the 100 resampling of validation sets within each water treatment (Figure 4) was likely a consequence of the small number of leaves in each validation set. An was poorly predicted with spectrometers alone regardless of the water treatment considered for the validation (Figure 4a, Supplemental Figure S7). In contrast, Li600 measurements allowed a good prediction of An, without much influence of water treatments, with median R² higher than 0.72 and median RMSE lower than 1.62 µmol m-2 s-1 across all validation sets (Figure 4). Neither the spectrometers nor the Li600 alone could reliably predict Ѱ (Figure 4).

**Figure 4.**
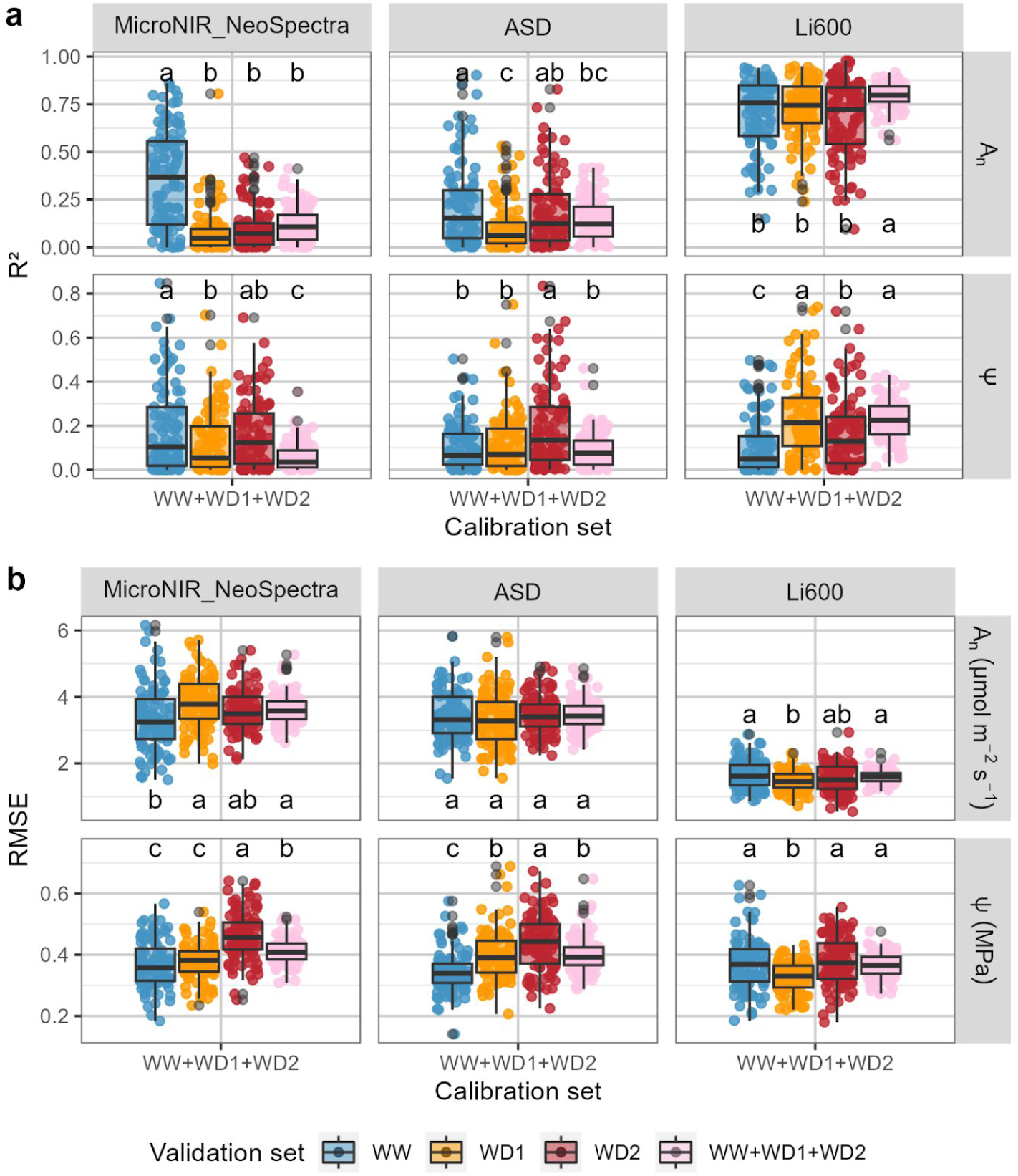
Prediction quality of PLSR models for An and Ѱ depending on the validation set. Models were calibrated using the WW+WD1+WD2 calibration set (Ddataset J in Table 1) and the MicroNIR_NeoSpectra, ASD and Li600 devices. a. corresponds to the R² and b. corresponds to the RMSE in the validation set. Each point in the boxplot represents one of the 100 iterations of the validation (as detailed in Table 1). The blue dots and boxes correspond to the well-watered treatment (WW); orange dots and boxes to the moderate water deficit treatment (WD1); red dots and boxes to the severe water deficit treatment (WD2); and pink dots and boxes to all WW, WD1 and WD2 treatments. A_n_: Net CO2 assimilation; Ѱ: Leaf water potential.

PLSR models built with Li600 measurements displayed a variable performance in predicting WUE_intr_ and WUE_inst_ depending on the validation set. The validation set that included the three water treatments (WW+WD1+WD2) showed intermediate prediction quality with median R² values of 0.52 and 0.46 for predicting WUE_intr_ and WUE_inst_, respectively (Figure 5a). If each treatment was considered separetely in the validation set, the prediction was higher for the WD1 water treatment, followed by intermediate performance under WW and finally poor performance under WD2 (Figure 5). These results indicated that PLSR models built with the Li600 device could not be considered accurate enough to predict WUE_intr_ and WUE_inst_ for leaves subjected to WD2 treatment. On the other hand, spectrometers performed poorly in predicting WUE_intr_ (Supplemental Figure S8) while MicroNIR and MicroNIR_NeoSpectra provided more relevant measurements to predict WUE_inst_ under WD1, with a median R² close to 0.53 (Supplemental Figure S8a).

**Figure 5.**
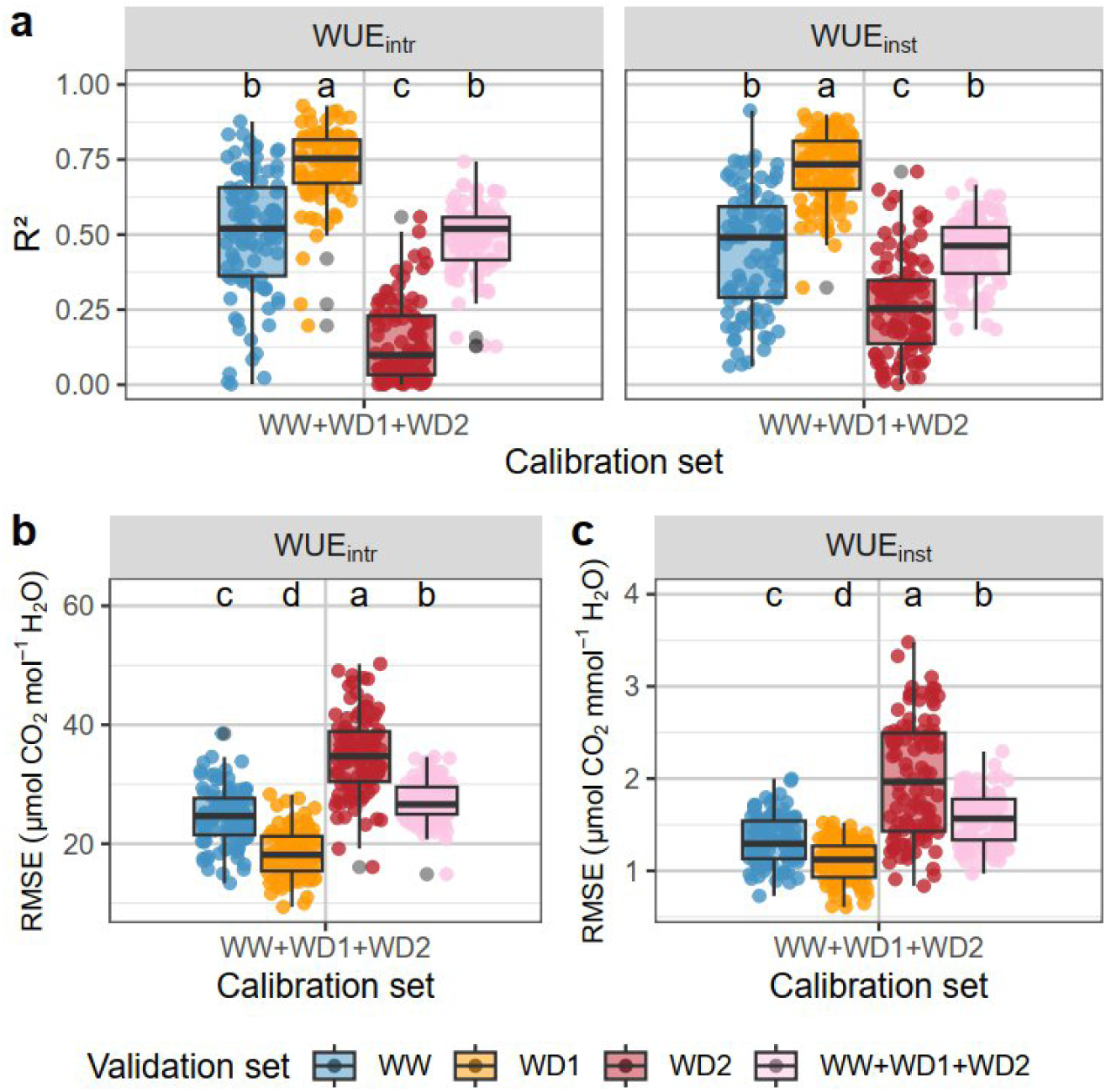
Prediction quality of PLSR models for WUE_intr_ and WUE_inst_ depending on the validation set. Models were calibrated with the WW+WD1+WD2 calibration set (Dataset J in Table 1) with variables from the Li600 device for intrinsic water use efficiency (WUE_intr_) and instantaneous water use efficiency (WUE_inst_) (as the ratio of Li6800 An and E or gsw measurements). a. corresponds to the R², b. corresponds to the RMSE expressed in µmol CO_2_ mol^−1^ H2O for WUE_intr_ and c corresponds to the RMSE expressed in µmol CO_2_ mmol^−1^ H_2_O for WUE_inst_. Each point in the boxplot corresponds to one of the 100 iterations of the validation (as detailed in Table 1). The blue dots and boxes correspond to the well-watered treatment (WW); orange dots and boxes to the moderate water deficit treatment (WD1); and red dots and boxes to the severe water deficit treatment (WD2).

### Prediction of LMA, A_n_, WUE_intr_ and WUE_inst_ using high-throughput measurements

We retained the PLSR models with a median R² higher than 0.55 in at least one validation set to make prediction within the entire panel while being trained with low throughput data (Dataset K in Table 1). The prediction models for LMA used between 3 and 12 latent variables, had R²cv higher than 0.76 and RMSE_cv_ lower than 0.4 mg cm-², irrespectively of the water treatment (Supplemental Figure S10a). Ten latent variables were retained to predict An using the Li600 device (Table 2). The corresponding model displayed good performance accross all water treatments with R²_cv_ and RMSE_cv_ of 0.8 and 1.59 µmol m-2 s-1 respectively. WUE_intr_ and WUE_inst_ were predicted with an R²_cv_ of 0.47 and 0.44, across all water treatment. In agreement with previous results, when considering leaves from WW and WD2 water treatments R^2^ did not exceed 0.53 whereas the R²_cv_ for WD1 leaves was above 0.73 (Supplemental Figure S10c, Supplemental Figure S10d). Therefore, models built with Li600 data were used to predict WUE_intr_ and WUE_inst_ for leaves under WD1 treatment only.

**Table 2.**
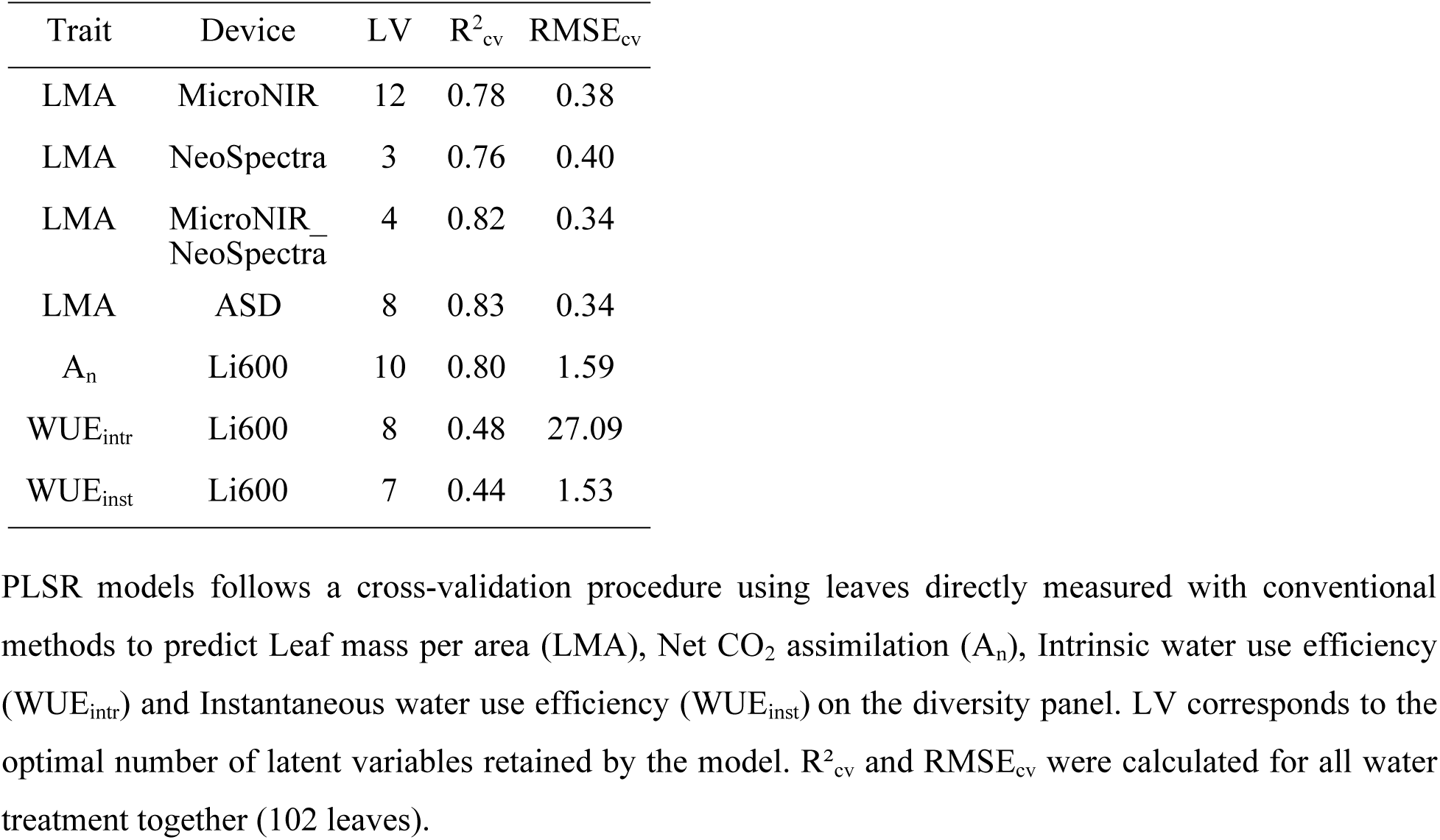
PLSR criteria obtained for predictive models (Dataset K inTable 1).

| Trait | Device | LV | $R^2_{cv}$ | $RMSE_{cv}$ |
| --- | --- | --- | --- | --- |
| LMA | MicroNIR | 12 | 0.78 | 0.38 |
| LMA | NeoSpectra | 3 | 0.76 | 0.40 |
| LMA | MicroNIR_<br>NeoSpectra | 4 | 0.82 | 0.34 |
| LMA | ASD | 8 | 0.83 | 0.34 |
| $A_n$ | Li600 | 10 | 0.80 | 1.59 |
| $WUE_{intr}$ | Li600 | 8 | 0.48 | 27.09 |
| $WUE_{inst}$ | Li600 | 7 | 0.44 | 1.53 |

PLSR models follows a cross-validation procedure using leaves directly measured with conventional methods to predict Leaf mass per area (LMA), Net $\text{CO}_2$ assimilation ( $A_n$ ), Intrinsic water use efficiency ( $WUE_{intr}$ ) and Instantaneous water use efficiency ( $WUE_{inst}$ ) on the diversity panel. LV corresponds to the optimal number of latent variables retained by the model. $R^2_{cv}$ and $RMSE_{cv}$ were calculated for all water treatment together (102 leaves).

The heritability of measured LMA across the entire diversity panel ranged from 0.28 to 0.43 depending on the water treatment (Figure 6a). The prediction of LMA using spectrometers provided very similar heritability estimates ranging from 0.20 to 0.42. We also estimated genetic values from LMA predictions which showed high correlations (r ≥ 0.9) with the genetic values estimated with the observed phenotypes, whatever the water treatment considered (Figure 6c). Predicted A_n_, WUE_intr_ and WUE_inst_ had heritability ranging from 0.16 to 0.5, depending on the water treatment (Figure 6b).

**Figure 6.**
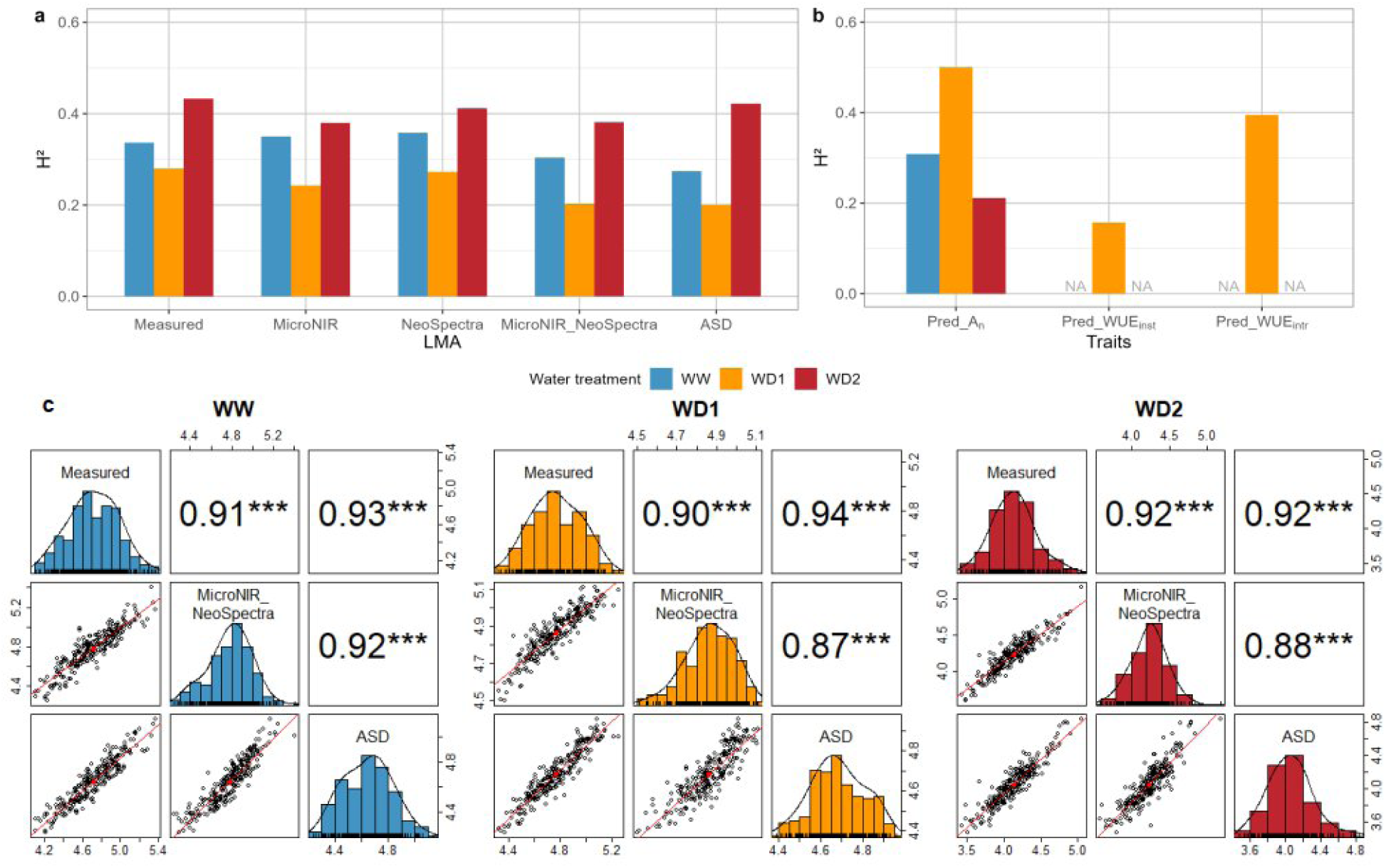
Heritability of observed and predicted leaf traits under different water treatments. a. Corresponds to the broad-sense heritability estimates for measured and predicted leaf mass per area (LMA) according to the spectrometer used as a predictor; b. corresponds to the heritability estimates for predicted net CO_2_ assimilation (Pred_A_n_), predicted instantaneous water use efficiency (Pred_WUE_inst_), predicted intrinsic water use efficiency (Pred_WUE_intr_); c. corresponds to the distributions and correlations between genotypic values of measured LMA versus genotypic values of predicted LMA according to the spectrometer used as a predictor (MicroNIR_NeoSpectra or ASD). Blue bars represent the well-watered treatment (WW); yellow bars represent the moderate water-deficit treatment (WD1); red bars represent the severe water deficit treatment (WD2). NA indicates that the trait was not predicted under the corresponding water treatment. ***: p-value ≤ 0.001.

## DISCUSSION

In this study, we explored the performance of models built with high-throughput devices to predict ecophysiological and morphological leaf traits across a large genetic diversity panel of *Vitis vinifera* varieties. Using a combination of large datasets collected on thousands of grapevine plants across different experiments and conditions, we were able to evaluate the robustness of predictive models with respect to the experimental conditions. Based on an analytical framework with dedicated and repeated calibration and validation data sets, we found that the prediction accuracy within the validation set containing leaves of plants grown under severe water deficit was generally lower than that for other water treatments. We also found that all devices (NIR spectrometers or poro-fluorometer) were efficient in predicting at least one trait. Finally, we found that the leaf traits predicted with NIRS and poro-fluorometry data captured a substantial part of the genetic variation which paves the way towards their use in genetic studies.

### High-throughput devices are complementary to predict a range of morphological and physiological traits

For too small datasets or too complex traits, repeated double cross-validation is more efficient to obtain an accurate estimation of the prediction error compared to a simple cross-validation procedure (Filzmoser et al., 2009). In our analyses, we performed 100 resampling iterations for evaluating the prediction of A_n_, Ѱ, WUE_intr_ and WUE_inst_ and observed that model R² was highly dependent on the iteration. While previous studies most often only made one iteration for cross-validating PLSR models (Van Wyngaard et al., 2021), our results underline the interest of repeating the cross-validation procedure to avoid bias in model evaluation.

Spectrometers resulted in better performances than Li600 for the prediction of LMA, WC_f_ and WQ while the Li600 was more efficient for the prediction of A_n_, WUE_intr_ and WUE_inst_. The laboratory spectrometer (ASD) was found as the most accurate to predict LMA. Absorption bands between 1670-1830 nm and 2000-2200 nm, covered by this device, were previously reported as relevant for predicting leaf dry matter related traits like protein, lignin and cellulose contents (Curran, 1989; Kokaly, 2001; Kawamura et al., 2008; Cheng et al., 2011). The difference of predictive performance between the laboratory spectrometer and the portable spectrometers could also be due to the fact that the ASD device were collected on dried leaves while spectra from MicroNIR and NeoSpectra devices were collected *in situ*. It is known that the presence of water in samples can dominate the signal, potentially reducing the detection of smaller peaks related to other components such as nitrogen (Masemola & Cho, 2019), protein, lignin and cellulose, which could be essential for predicting LMA. LMA and WC_f_ were negatively correlated, likely because leaves with a low LMA tend to have a high water content (Poorter et al., 2009), but we observed that the devices yielding the best PLSR models for predicting these traits differed. Indeed, in that case contrary to LMA, water-related traits (WC_f_ and WQ) were better predicted *in situ* using portable spectrometers. Moreover, water-related traits were better predicted using the combined wavelengths of MicroNIR and NeoSpectra than with either device alone. This is likely because their combined spectral range includes the two main water absorption bands around 1400–1440 nm and 1900–1950 nm (De Bei et al., 2011).

The poro-fluorometer device was found to provide good prediction of A_n_. To our knowledge. This aligns with previous studies showing that A_n_ correlates with the fluorescence decline ratio under steady-state conditions with saturated light (Lichtenthaler et al., 2005). Additionally, A_n_ was well predicted when combining the electron transport rate, also derived from leaf fluorescence measurements, with the leaf-to-air temperature difference, used as a proxy for stomatal closure (Han et al., 2022; Losciale et al., 2015; Coupel-Ledru et al., 2019). In our study, spectrometers could not accurately predict A_n_, in contrast to a previous study on maize (Cotrozzi et al., 2020). However, at least 3 spectra from the same side of a leaf were used in the maize experiment, while in our study, two spectra (one on each side) were taken from the same spatial area of a given leaf.

We found that the Li600 device was able to predict WUE_intr_ and WUE_inst_ for plants under a moderate water deficit. In contrast, spectral data generally failed to provide reliable predictions, except for WUE_inst_, which could be predicted using MicroNIR and the MicroNIR-NeoSpectra combination under moderate water deficit. With a larger sample size, (Cotrozzi et al., 2020) reported fairly good predictive performance of NIRS models for WUE_intr_ and WUE_inst_ in maize.

The leaf water potential is a physiological trait commonly used as an indicator of soil water deficit intensity (Simonneau et al., 2017; De Bei et al., 2011; Girona et al., 2006). In our study, the leaf water potential was poorly predicted, regardless of the device used. Experiments on maize have shown that the leaf water potential could be predicted by NIRS with a R²_cv_ of 0.63 on both greenhouse and field samples (Cotrozzi et al., 2020). Field experiments on grapevines have shown the interest of NIRS for the prediction of leaf water potential on Syrah and Cabernet Sauvignon (De Bei et al., 2011). However, in these studies, stem water potential was better predicted than leaf water potential perhaps because spectra, taken at a single spot on the leaf, are not representative of the spatial variability in stomatal conductance (De Bei et al., 2011; Albasha et al., 2019).

### The robustness of PLSR models depends on the traits and the plant growing conditions

LMA has been widely used across species and locations to investigate the robustness of predictive models when plants were subjected to different environmental conditions (Serbin et al., 2019; Ji et al., 2024). In these studies, a considerable decrease in model accuracy was found when a trained PLSR model was applied to a validation set containing other sites or experiments. Here, we consistently highlighted a degradation of outdoors prediction using a greenhouse calibration set, similar to recent reports on maize (Cotrozzi et al., 2020). In present work, we further aimed at deciphering the effect of the watering conditions on the PLSR models prediction accuracy. The prediction of ecophysiological traits might indeed be affected by soil water availability. While most published studies pooled together data collected in different growing conditions in the calibration and validation datasets (De Bei et al., 2011; Ryckewaert et al., 2022b), here we tested both the effects of calibrating PLSR models per water treatment, and of validating them under different water treatments. Our results highlighted that the composition of the validation dataset was more influential than the calibration one. Validating PLSR models per water treatment prevents an artificial inflation of R^2^ value caused by the inclusion of multiple water treatments which in turn increases the trait variance. This likely led to more accurate PLSR model evaluations which highlighted changes in prediction accuracy for LMA, WC_f_, WUE_intr_ and WUE_inst_ depending on the water treatments. Such variation in R^2^ is primarily due to the reduction in trait variance as underlined by the smaller differences observed for RMSE across water treatments.

### PLSR models allow capturing the genetic variability of leaf traits

High correlations between predicted and measured values do not necessarily imply an accurate prediction of the genetic variability of the traits, especially when the calibration set includes multiple experimental conditions. In such cases, the model might have been trained to predict differences between experimental treatments rather than between plant-related variation within a given treatment.

Our results demonstrated that the broad sense heritability of predicted LMA was comparable to that of measured LMA, underlying, at least for this trait, that predicted values were as efficient as measured values for capturing the genetic variability within each water treatment. Similar findings were reported for chlorophyll content, nitrogen content and specific leaf area, predicted using hyperspectral reflectance data in maize grown under low and high nitrogen conditions (Grzybowski et al., 2021). The heritability of predicted A_n_, WUE_intr_ and WUE_inst_ depended on the water treatment with higher heritabilities for A_n_ and WUE_intr_ under moderate water deficit compared to WUE_inst_ and other water treatments. One reason to explain these results could be that our study was based on a single leaf per plant, even though the choice of the leaf was homogeneous between plants and chosen as well-exposed and fully expanded. Increasing the number of measurements is likely to increase trait heritability (Jablonszky & Garamszegi, 2024), but this will also extend the time needed to performed the measurements, which may not be appropriate in a high-throughput phenotyping strategy. To overcome this problem, other methodologies based on contactless devices could be tested, such as hyper-spectral imaging (Zhang et al., 2023). Thermal imaging could also be used to assess leaves and canopy temperature as proxies of water stress responses (Wen et al., 2023). Architectural and functional traits have successfully been characterized using LiDAR, multispectral and thermal imaging in a large core-collection of apple trees (Coupel-Ledru et al., 2019), highlighting a strong genotypic effect on all traits, with high broad-sense heritability. A significant effect of the water treatment was also detected on most of the traits. However, when traits exhibit high spatial variability at fine scale, the accuracy of phenotyping using imaging sensors is related to the image resolution. Airborne imaging may also be affected by background noise and atmospheric scattering, which can alter the light absorption features of pigments (Tattaris et al., 2016). Nevertheless, these high-throughput methods are highly effective for phenotyping large numbers of plants and multiple environments in a short time, thus reducing the error associated with the plant circadian cycle. These phenotyping methods open prospects for further analyses of the genetic determinism of ecophysiological traits as recently shown in apple trees (Coupel-Ledru et al., 2022).

## CONCLUSION

Our findings demonstrate the strengths of high-throughput devices for predicting ecophysiological traits not directly accessible at the same throughput in a diversity panel of grapevine under varying experimental conditions. We have highlighted the importance of validating PLSR models for each experimental condition and not being restricted to a single calibration/validation set. This approach can be applied to other abiotic stresses, other populations or even other species. This would require sharing high-throughput dataset collected on multiple species, growth conditions and environments to increase predictive models robustness so that they can be applied more easily to a wide range of study cases. High-throughput phenotyping strategies can be integrated with imaging-based phenotyping or 3D modeling to improve the assessment of inter-plant variability and further enhance our understanding of plant responses to environmental stresses through genetic analyses. Overall our results open promising avenues for further deploying genome-wide association studies on traits predicted from high-throughput measurements.

## Abbreviations

A_n_: net photosynthesis
DW: dry weight
E: transpiration
FW: fresh weight
g_sw_: stomatal conductance for water vapor
IRGA: infrared gas-exchange analyzer
LMA: leaf mass area
LOO: leave-one-out
NIRS: near-infrared spectroscopy
NRMSE: normalized root mean squared error
PLSR: partial least square regression
PPFD: photosynthetically photon flux density
RMSE: root mean squared error
WC_f_: leaf water content
WD1: moderate Water Deficit
WD2: severe Water Deficit
WQ: leaf water quantity
WUE_inst_: instantaneous water use efficiency
WUE_intr_: intrinsic water use efficiency ;
WW: Well-Watered
Ѱ: leaf water potential.

## ACKNOWLEDGMENTS

We are grateful to the INRAE “Domaine de Vassal” Grape Germplasm Repository for collection maintenance. We thank Freddy Gavanon and Mélyne Falcon for the plant management and the preparation of the greenhouse experiment. We thank Gaëlle Rolland for her help in the leaf phenotyping. We thank Martin Ecarnot and Frederic Compan for the access to the laboratory spectrometer ASD. We thank the ETAP (LEPSE), M3P (LEPSE) and DAAV (AGAP Institut) team members for providing help during the preparation and phenotyping of the grapevine panel. We thank Laurent Torregrosa for administration of ANR G2WAS project and Dominique This and Roberto Bacilieri for administration of Défi clé Vinid’Occ PlastiVigne.

## SUPPLEMENTAL MATERIAL

Supplemental material includes 3 supplemental tables ans 9 supplemental figures.

## OPTIONAL SECTIONS

### Data availability

The datasets supporting the conclusions of this article are available in the Recherche Data Gouv repository at https://doi.org/10.57745/WVAPOL. The scripts for prediction analyses are publicly available in GitLab: https://forgemia.inra.fr/eva.coindre/robustness_ht_prediction_ecophysiological_traits.

### Author contributions

AC-L, VS designed the project and managed the outdoors experiment. TS, AD, AC-L, VS managed the greenhouse experiment in the frame of the G²WAS project. ML and LC-B managed the PhenoArch platform. RB, LC, VF, TL, VB, EC, TS, AC-L and VS collected data. RB, LC and EC curated data. LC and VF performed preliminary analyses. EC analyzed data with input from MR, BP, AC-L and VS. AC-L, VS and BP supervised research. EC, AC-L, BP and VS wrote the manuscript with input from all authors.

### Funding

This research was supported by the ANR (French Research Agency) through the G²WAS project (ANR-19-CE20-0024) and the #Digitag project (ANR-16-CONV-0004). PhenoArch platform is funded by the project ANR-11-INBS-0012 (PHENOME-EMPHASIS). This work was publicly funded through the Occitanie Region’s program “Défi clé Vinid’Occ” run by the University of Montpellier (PhD grant to EC within the project PlastiVigne).

## REFERENCES

Albasha, R., Fournier, C., Pradal, C., Chelle, M., Prieto, J. A., Louarn, G., Simonneau, T., & Lebon, E. (2019). HydroShoot : A functional-structural plant model for simulating hydraulic structure, gas and energy exchange dynamics of complex plant canopies under water deficit—application to grapevine ( *Vitis vinifera* ). In Silico Plants, 1(1), diz007. 10.1093/insilicoplants/diz007

Barnes, R. J., Dhanoa, M. S., & Lister, S. J. (1989). Standard Normal Variate Transformation and De-Trending of Near-Infrared Diffuse Reflectance Spectra. Applied Spectroscopy, 43(5), 772-777. 10.1366/0003702894202201

Barthès, B. G., Kouakoua, E., Clairotte, M., Lallemand, J., Chapuis-Lardy, L., Rabenarivo, M., & Roussel, S. (2019). Performance comparison between a miniaturized and a conventional near infrared reflectance (NIR) spectrometer for characterizing soil carbon and nitrogen. Geoderma, 338, 422-429. 10.1016/j.geoderma.2018.12.031

Bates, D., Mächler, M., Bolker, B., & Walker, S. (2015). Fitting Linear Mixed-Effects Models Using **lme4**. Journal of Statistical Software, 67(1). 10.18637/jss.v067.i01

Bhattacharya, A. (2021). Effect of Soil Water Deficit on Growth and Development of Plants : A Review. In A. Bhattacharya, Soil Water Deficit and Physiological Issues in Plants (p. 393-488). Springer Singapore. 10.1007/978-981-33-6276-5_5

Bota, B. J., Flexas, J., & Medrano, H. (2001). Genetic variability of photosynthesis and water use in Balearic grapevine cultivars. Annals of Applied Biology, 138(3), 353-361. 10.1111/j.1744-7348.2001.tb00120.x

Brandolini-Bunlon, M., Jaillais, B., Roger, J.-M., & Lesnoff, M. (2023). rchemo : Dimension Reduction, Regression and Discrimination for Chemometrics (p. 0.1–3) [Jeu de données]. 10.32614/CRAN.package.rchemo

Cabrera-Bosquet, L., Crossa, J., Von Zitzewitz, J., Serret, M. D., & Luis Araus, J. (2012). High-throughput Phenotyping and Genomic Selection : The Frontiers of Crop Breeding Converge^F^. Journal of Integrative Plant Biology, 54(5), 312-320. 10.1111/j.1744-7909.2012.01116.x

Cabrera-Bosquet, L., Fournier, C., Brichet, N., Welcker, C., Suard, B., & Tardieu, F. (2016). High-throughput estimation of incident light, light interception and radiation-use efficiency of thousands of plants in a phenotyping platform. New Phytologist, 212(1), 269-281. 10.1111/nph.14027

Calvin, K., Dasgupta, D., Krinner, G., Mukherji, A., Thorne, P. W., Trisos, C., Romero, J., Aldunce, P., Barrett, K., Blanco, G., Cheung, W. W. L., Connors, S., Denton, F., Diongue-Niang, A., Dodman, D., Garschagen, M., Geden, O., Hayward, B., Jones, C., … Péan, C. (2023). IPCC, 2023 : Climate Change 2023: Synthesis Report. Contribution of Working Groups I, II and III to the Sixth Assessment Report of the Intergovernmental Panel on Climate Change *[Core Writing Team,* H. Lee and J. Romero *(eds.)]*. IPCC, Geneva, Switzerland. (First). Intergovernmental Panel on Climate Change (IPCC). 10.59327/IPCC/AR6-9789291691647

Cheng, T., Rivard, B., & Sánchez-Azofeifa, A. (2011). Spectroscopic determination of leaf water content using continuous wavelet analysis. Remote Sensing of Environment, 115(2), 659-670. 10.1016/j.rse.2010.11.001

Cotrozzi, L., Peron, R., Tuinstra, M. R., Mickelbart, M. V., & Couture, J. J. (2020). Spectral Phenotyping of Physiological and Anatomical Leaf Traits Related with Maize Water Status. Plant Physiology, 184(3), 1363-1377. 10.1104/pp.20.00577

Coupel-Ledru, A. (2021). Plant Water-Use Efficiency. In John Wiley & Sons, Ltd (Éd.), *eLS* (1^re^ éd., p. 1-8). Wiley. 10.1002/9780470015902.a0027971

Coupel-Ledru, A., Lebon, É., Christophe, A., Doligez, A., Cabrera-Bosquet, L., Péchier, P., Hamard, P., This, P., & Simonneau, T. (2014). Genetic variation in a grapevine progeny (Vitis vinifera L. cvs Grenache×Syrah) reveals inconsistencies between maintenance of daytime leaf water potential and response of transpiration rate under drought. Journal of Experimental Botany, 65(21), 6205-6218. 10.1093/jxb/eru228

Coupel-Ledru, A., Pallas, B., Delalande, M., Boudon, F., Carrié, E., Martinez, S., Regnard, J.-L., & Costes, E. (2019). Multi-scale high-throughput phenotyping of apple architectural and functional traits in orchard reveals genotypic variability under contrasted watering regimes. Horticulture Research, 6(1), 52. 10.1038/s41438-019-0137-3

Coupel-Ledru, A., Pallas, B., Delalande, M., Segura, V., Guitton, B., Muranty, H., Durel, C., Regnard, J., & Costes, E. (2022). Tree architecture, light interception and water-use related traits are controlled by different genomic regions in an apple tree core collection. New Phytologist, 234(1), 209-226. 10.1111/nph.17960

Coupel-Ledru, A., Westgeest, A. J., Albasha, R., Millan, M., Pallas, B., Doligez, A., Flutre, T., Segura, V., This, P., Torregrosa, L., Simonneau, T., & Pantin, F. (2024). Clusters of grapevine genes for a burning world. New Phytologist, nph.19540. 10.1111/nph.19540

Curran, P. J. (1989). Remote sensing of foliar chemistry. Remote Sensing of Environment, 30(3), 271-278. 10.1016/0034-4257(89)90069-2

Damour, G., Simonneau, T., Cochard, H., & Urban, L. (2010). An overview of models of stomatal conductance at the leaf level : Models of stomatal conductance. Plant, Cell & Environment, no-no. 10.1111/j.1365-3040.2010.02181.x

Das, B., Sahoo, R. N., Pargal, S., Krishna, G., Verma, R., Viswanathan, C., Sehgal, V. K., & Gupta, V. K. (2021). Evaluation of different water absorption bands, indices and multivariate models for water-deficit stress monitoring in rice using visible-near infrared spectroscopy. Spectrochimica Acta Part A: Molecular and Biomolecular Spectroscopy, 247, 119104. 10.1016/j.saa.2020.119104

De Bei, R., Cozzolino, D., Sullivan, W., Cynkar, W., Fuentes, S., Dambergs, R., Pech, J., & Tyerman, S. (2011). Non-destructive measurement of grapevine water potential using near infrared spectroscopy : Measure of grapevine water potential using NIR. Australian Journal of Grape and Wine Research, 17(1), 62-71. 10.1111/j.1755-0238.2010.00117.x

De Bei, R., Fuentes, S., Sullivan, W., Edwards, E. J., Tyerman, S., & Cozzolino, D. (2017). Rapid measurement of total non-structural carbohydrate concentration in grapevine trunk and leaf tissues using near infrared spectroscopy. Computers and Electronics in Agriculture, 136, 176-183. 10.1016/j.compag.2017.03.007

Ecarnot, M., Compan, F., & Roumet, P. (2013). Assessing leaf nitrogen content and leaf mass per unit area of wheat in the field throughout plant cycle with a portable spectrometer. Field Crops Research, 140, 44-50. 10.1016/j.fcr.2012.10.013

Ely, K. S., Burnett, A. C., Lieberman-Cribbin, W., Serbin, S. P., & Rogers, A. (2019). Spectroscopy can predict key leaf traits associated with source–sink balance and carbon–nitrogen status. Journal of Experimental Botany, 70(6), 1789-1799. 10.1093/jxb/erz061

Fernández-Calleja, M., Monteagudo, A., Casas, A. M., Boutin, C., Pin, P. A., Morales, F., & Igartua, E. (2020). Rapid On-Site Phenotyping via Field Fluorimeter Detects Differences in Photosynthetic Performance in a Hybrid—Parent Barley Germplasm Set. Sensors, 20(5), 1486. 10.3390/s20051486

Filzmoser, P., Liebmann, B., & Varmuza, K. (2009). Repeated double cross validation. Journal of Chemometrics, 23(4), 160-171. 10.1002/cem.1225

Flutre, T. & Chabrault. (2019). timflutre/rutilstimflutre : V0.170.0 (Version v0.170.0) [Logiciel]. Zenodo. 10.5281/ZENODO.3580267

Foley, W. J., McIlwee, A., Lawler, I., Aragones, L., Woolnough, A. P., & Berding, N. (1998). Ecological applications of near infrared reflectance spectroscopy—A tool for rapid, cost-effective prediction of the composition of plant and animal tissues and aspects of animal performance. Oecologia, 116(3), 293-305. 10.1007/s004420050591

Ge, Y., Atefi, A., Zhang, H., Miao, C., Ramamurthy, R. K., Sigmon, B., Yang, J., & Schnable, J. C. (2019). High-throughput analysis of leaf physiological and chemical traits with VIS–NIR–SWIR spectroscopy : A case study with a maize diversity panel. Plant Methods, 15(1), 66. 10.1186/s13007-019-0450-8

Girona, J., Mata, M., Del Campo, J., Arbonés, A., Bartra, E., & Marsal, J. (2006). The use of midday leaf water potential for scheduling deficit irrigation in vineyards. Irrigation Science, 24(2), 115-127. 10.1007/s00271-005-0015-7

Grzybowski, M., Wijewardane, N. K., Atefi, A., Ge, Y., & Schnable, J. C. (2021). Hyperspectral reflectance-based phenotyping for quantitative genetics in crops : Progress and challenges. Plant Communications, 2(4), 100209. 10.1016/j.xplc.2021.100209

Han, J., Chang, C. Y., Gu, L., Zhang, Y., Meeker, E. W., Magney, T. S., Walker, A. P., Wen, J., Kira, O., McNaull, S., & Sun, Y. (2022). The physiological basis for estimating photosynthesis from Chl *a* fluorescence. New Phytologist, 234(4), 1206-1219. 10.1111/nph.18045

Jablonszky, M., & Garamszegi, L. Z. (2024). The effect of repeated measurements and within-individual variance on the estimation of heritability : A simulation study. Behavioral Ecology and Sociobiology, 78(2), 18. 10.1007/s00265-024-03435-w

Ji, F., Li, F., Hao, D., Shiklomanov, A. N., Yang, X., Townsend, P. A., Dashti, H., Nakaji, T., Kovach, K. R., Liu, H., Luo, M., & Chen, M. (2024). Unveiling the transferability of PLSR models for leaf trait estimation : Lessons from a comprehensive analysis with a novel global dataset. New Phytologist, 243(1), 111-131. 10.1111/nph.19807

Kawamura, K., Watanabe, N., Sakanoue, S., & Inoue, Y. (2008). Estimating forage biomass and quality in a mixed sown pasture based on partial least squares regression with waveband selection. Grassland Science, 54(3), 131-145. 10.1111/j.1744-697X.2008.00116.x

Kokaly, R. F. (2001). Investigating a Physical Basis for Spectroscopic Estimates of Leaf Nitrogen Concentration. Remote Sensing of Environment, 75(2), 153-161. 10.1016/S0034-4257(00)00163-2

Lichtenthaler, H. K., Buschmann, C., & Knapp, M. (2005). How to correctly determine the different chlorophyll fluorescence parameters and the chlorophyll fluorescence decrease ratio R_Fd_ of leaves with the PAM fluorometer. Photosynthetica, 43(3), 379-393. 10.1007/s11099-005-0062-6

Liland, K. H., Mevik, B.-H., & Wehrens, R. (1999). pls : Partial Least Squares and Principal Component Regression (p. 2.8-5) [Jeu de données]. 10.32614/CRAN.package.pls

Losciale, P., Manfrini, L., Morandi, B., Pierpaoli, E., Zibordi, M., Stellacci, A. M., Salvati, L., & Corelli Grappadelli, L. (2015). A multivariate approach for assessing leaf photo-assimilation performance using the I_PL_ index. Physiologia Plantarum, 154(4), 609-620. 10.1111/ppl.12328

Masemola, C., & Cho, M. A. (2019). Estimating leaf nitrogen concentration from similarities in fresh and dry leaf spectral bands using a model population analysis framework. International Journal of Remote Sensing, 40(17), 6841-6860. 10.1080/01431161.2019.1597300

Murchie, E. H., & Lawson, T. (2013). Chlorophyll fluorescence analysis : A guide to good practice and understanding some new applications. Journal of Experimental Botany, 64(13), 3983-3998. 10.1093/jxb/ert208

Nicolas, S. D., Péros, J.-P., Lacombe, T., Launay, A., Le Paslier, M.-C., Bérard, A., Mangin, B., Valière, S., Martins, F., Le Cunff, L., Laucou, V., Bacilieri, R., Dereeper, A., Chatelet, P., This, P., & Doligez, A. (2016). Genetic diversity, linkage disequilibrium and power of a large grapevine (Vitis vinifera L) diversity panel newly designed for association studies. BMC Plant Biology, 16(1), 74. 10.1186/s12870-016-0754-z

Petisco, C., García-Criado, B., Mediavilla, S., Vázquez De Aldana, B. R., Zabalgogeazcoa, I., & García-Ciudad, A. (2006). Near-infrared reflectance spectroscopy as a fast and non-destructive tool to predict foliar organic constituents of several woody species. Analytical and Bioanalytical Chemistry, 386(6), 1823-1833. 10.1007/s00216-006-0816-4

Poorter, H., Niinemets, Ü., Poorter, L., Wright, I. J., & Villar, R. (2009). Causes and consequences of variation in leaf mass per area (LMA) : A meta-analysis. New Phytologist, 182(3), 565-588. 10.1111/j.1469-8137.2009.02830.x

Prieto, J. A., Lebon, É., & Ojeda, H. (2010). Stomatal behavior of different grapevine cultivars in response to soil water status and air water vapor pressure deficit. OENO One, 44(1), 9. 10.20870/oeno-one.2010.44.1.1459

R Core Team. (2023). R: A Language and Environment for Statistical Computing. R Foundation for Statistical Computing. https://www.R-project.org/

Rapaport, T., Hochberg, U., Shoshany, M., Karnieli, A., & Rachmilevitch, S. (2015). Combining leaf physiology, hyperspectral imaging and partial least squares-regression (PLS-R) for grapevine water status assessment. ISPRS Journal of Photogrammetry and Remote Sensing, 109, 88-97. 10.1016/j.isprsjprs.2015.09.003

Ryckewaert, M., Héran, D., Simonneau, T., Abdelghafour, F., Boulord, R., Saurin, N., Moura, D., Mas-Garcia, S., & Bendoula, R. (2022a). Physiological variable predictions using VIS–NIR spectroscopy for water stress detection on grapevine : Interest in combining climate data using multiblock method. Computers and Electronics in Agriculture, 197, 106973. 10.1016/j.compag.2022.106973

Ryckewaert, M., Héran, D., Simonneau, T., Abdelghafour, F., Boulord, R., Saurin, N., Moura, D., Mas-Garcia, S., & Bendoula, R. (2022b). Physiological variable predictions using VIS–NIR spectroscopy for water stress detection on grapevine : Interest in combining climate data using multiblock method. Computers and Electronics in Agriculture, 197, 106973. 10.1016/j.compag.2022.106973

Savitzky, Abraham., & Golay, M. J. E. (1964). Smoothing and Differentiation of Data by Simplified Least Squares Procedures. Analytical Chemistry, 36(8), 1627-1639. 10.1021/ac60214a047

Semia Cherif, Doblas-Miranda, E., Lionello, P., Borrego, C., Giorgi, F., Iglesias, A., Sihem Jebari, Mahmoudi, E., Moriondo, M., Pringault, O., Rilov, G., Somot, S., Tsikliras, A., Vilà, M., & Zittis, G. (2020). First Mediterranean Assessment Report - Chapter 2 : Drivers of Change. Zenodo. 10.5281/ZENODO.7100601

Serbin, S. P., Wu, J., Ely, K. S., Kruger, E. L., Townsend, P. A., Meng, R., Wolfe, B. T., Chlus, A., Wang, Z., & Rogers, A. (2019). From the Arctic to the tropics : Multibiome prediction of leaf mass per area using leaf reflectance. New Phytologist, 224(4), 1557-1568. 10.1111/nph.16123

signal developers. (2023). signal : Signal processing. https://r-forge.r-project.org/projects/signal/

Silva-Perez, V., Molero, G., Serbin, S. P., Condon, A. G., Reynolds, M. P., Furbank, R. T., & Evans, J. R. (2018). Hyperspectral reflectance as a tool to measure biochemical and physiological traits in wheat. Journal of Experimental Botany, 69(3), 483-496. 10.1093/jxb/erx421

Simonneau, T., Lebon, E., Coupel-Ledru, A., Marguerit, E., Rossdeutsch, L., & Ollat, N. (2017). Adapting plant material to face water stress in vineyards : Which physiological targets for an optimal control of plant water status? OENO One, 51(2), 167. 10.20870/oeno-one.2016.0.0.1870

Tattaris, M., Reynolds, M. P., & Chapman, S. C. (2016). A Direct Comparison of Remote Sensing Approaches for High-Throughput Phenotyping in Plant Breeding. Frontiers in Plant Science, 7. 10.3389/fpls.2016.01131

Tomás, M., Medrano, H., Escalona, J. M., Martorell, S., Pou, A., Ribas-Carbó, M., & Flexas, J. (2014). Variability of water use efficiency in grapevines. Environmental and Experimental Botany, 103, 148-157. 10.1016/j.envexpbot.2013.09.003

Trenti, M., Lorenzi, S., Bianchedi, P. L., Grossi, D., Failla, O., Grando, M. S., & Emanuelli, F. (2021). Candidate genes and SNPs associated with stomatal conductance under drought stress in Vitis. BMC Plant Biology, 21(1), 7. 10.1186/s12870-020-02739-z

Van Leeuwen, C., Sgubin, G., Bois, B., Ollat, N., Swingedouw, D., Zito, S., & Gambetta, G. A. (2024). Climate change impacts and adaptations of wine production. Nature Reviews Earth & Environment, 5(4), 258-275. 10.1038/s43017-024-00521-5

Van Wyngaard, E., Blancquaert, E., Nieuwoudt, H., & Aleixandre-Tudo, J. L. (2021). Infrared Spectroscopy and Chemometric Applications for the Qualitative and Quantitative Investigation of Grapevine Organs. Frontiers in Plant Science, 12, 723247. 10.3389/fpls.2021.723247

Wen, T., Li, J.-H., Wang, Q., Gao, Y.-Y., Hao, G.-F., & Song, B.-A. (2023). Thermal imaging : The digital eye facilitates high-throughput phenotyping traits of plant growth and stress responses. Science of The Total Environment, 899, 165626. 10.1016/j.scitotenv.2023.165626

Wold, S., Sjöström, M., & Eriksson, L. (2001). PLS-regression : A basic tool of chemometrics. Chemometrics and Intelligent Laboratory Systems, 58(2), 109-130. 10.1016/S0169-7439(01)00155-1

Workman Jr., J., & Weyer, L. (2012). Practical Guide and Spectral Atlas for Interpretive Near-Infrared Spectroscopy (0 éd.). CRC Press. 10.1201/b11894

Xiao, Q., Bai, X., Zhang, C., & He, Y. (2022). Advanced high-throughput plant phenotyping techniques for genome-wide association studies : A review. Journal of Advanced Research, 35, 215-230. 10.1016/j.jare.2021.05.002

Zhang, H., Wang, L., Jin, X., Bian, L., & Ge, Y. (2023). High-throughput phenotyping of plant leaf morphological, physiological, and biochemical traits on multiple scales using optical sensing. The Crop Journal, 11(5), 1303-1318. 10.1016/j.cj.2023.04.014

Zhu, C., Fu, X., Zhang, J., Qin, K., & Wu, C. (2022). Review of portable near infrared spectrometers : Current status and new techniques. Journal of Near Infrared Spectroscopy, 30(2), 51-66. 10.1177/09670335211030617

