## Supplementary materials for "Robustness of high-throughput prediction of leaf ecophysiological traits using near infrared spectroscopy and poro-fluorometry"

**Supplemental Table S1. Measured variables from the poro-fluorometer Li600 device.**

| Type of variable | Description | Label | Units |
|---|---|---|---|
| Porometry variables | Stomatal conductance | gsw | mol $m^{-2}$ $s^{-1}$ |
| | Transpiration rate | E_apparent | mol $m^{-2}$ $s^{-2}$ |
| | One-sided boundary layer conductance | gbw | mol $m^{-2}$ $s^{-2}$ |
| | Total conductance | gtw | mol $m^{-2}$ $s^{-2}$ |
| | Chamber vapor pressure | Vpcham | kPa |
| | Reference vapor pressure | Vpref | kPa |
| | Leaf vapor pressure | VPleaf | kPa |
| | Leaf vapor pressure deficit | VPDleaf | kPa |
| | Reference $H_2O$ mole fraction | H2O_r | mmol $mol^{-1}$ |
| | Sample $H_2O$ mole fraction | H2O_s | mmol $mol^{-1}$ |
| | Leaf $H_2O$ mole fraction | H2O_leaf | mmol $mol^{-1}$ |
| Fluorescence variables | Minimum fluorescence in light | Fs | NA |
| | Maximum fluorescence in light | Fm' | NA |
| | Quantum efficiency of photosystem 2 in light | PhiPS2 | NA |
| | Electron transport rate | ETR | µmol $m^{-2}$ $s^{-1}$ |
| Environmental variables | Sample relative humidity | rh_s | % |
| | Reference relative humidity | rh_r | % |
| | Block reference temperature | Tref | °C |
| | Leaf temperature | Tleaf | °C |
| | Atmospheric pressure | P_atm | kPa |
| | Ambient light | Qamb | µmol $m^{-2}$ $s^{-1}$ |

**Supplemental Table S2. Coefficient of variation for leaf traits directly measured with conventional methods.**

The coefficient of variation is calculated as $\frac{\sigma}{|\bar{x}|} \times 100$ with σ the standard deviation and $|\bar{x}|$ the absolute mean of the trait. HT = high- and LT = low-throughput measurements, exp. = experiment.

| Experiment | Variable name | Unit | Measuring method | HT or LT | Leaves number | Coefficient of variation | | |
|---|---|---|---|---|---|---|---|---|
| Outdoor exp. | Leaf Mass Area (LMA) | mg $cm^{-2}$ | Disk weighing | HT | 470 | 16.7 | | |
| | | | | | | **WW** | **WD1** | **WD2** |
| Greenhouse exp. | Leaf mass per area (LMA) | mg $cm^{-2}$ | Disk weighing | HT | 1410 | 21.4 | 20.5 | 21.4 |
| | | | | LT | 102 | 22.3 | 19 | 15.7 |
| | Water content ($WC_f$) | mg $H_2O$ $mg^{-1}$ fresh weight | Disk weighing | HT | 1410 | 4.7 | 4.3 | 3.7 |
| | Water quantity (WQ) | mg $H_2O$ | Disk weighing | HT | 1410 | 20 | 18.8 | 19.9 |
| | Net $CO_2$ assimilation ($A_n$) | µmol $m^{-2}$ $s^{-1}$ | Li6800 | LT | 102 | 35.9 | 42.4 | 65.4 |
| | Water potential (Ψ) | MPa | Pressure chamber | LT | 102 | 32.2 | 39.2 | 29.4 |
| | Intrinsic water use efficiency ($WUE_{intr}$) | µmol $CO_2$ $mol^{-1}$ $H_2O$ | Li6800 | LT | 102 | 42.7 | 38.8 | 32.1 |
| | Instantaneous water use efficiency ($WUE_{inst}$) | µmol $CO_2$ $mmol^{-1}$ $H_2O$ | Li6800 | LT | 102 | 44 | 44.9 | 46.6 |

**Supplemental Table S3. Student's t test on $R^2_{cv}$ for each device combination for each predicted trait (datasets A and B in Table 1)** : Leaf mass per area (LMA), Water content ($WC_f$), Water quantity (WQ), Net $CO_2$ assimilation ($A_n$), Leaf water potential (Ψ), Intrinsic water use efficiency ($WUE_{intr}$) and Instantaneous water use efficiency ($WUE_{inst}$). ns = p-value ≥ 0.05; * = p-value ≤ 0.05; ** = p-value ≤ 0.01; *** = p-value ≤ 0.001; **** = p-value ≤ 1e-04.

| Device1 | Device2 | LMA | $WC_f$ | WQ | $A_n$ | Ψ | $WUE_{intr}$ | $WUE_{inst}$ |
|---|---|---|---|---|---|---|---|---|
| ASD | Li600 | **** | **** | **** | **** | **** | **** | **** |
| ASD | MicroNIR | **** | **** | **** | **** | ns | **** | ns |
| ASD | MicroNIR_NeoSpectra | **** | **** | **** | **** | ns | **** | **** |
| ASD | NeoSpectra | **** | **** | **** | *** | *** | ns | **** |
| Li600 | MicroNIR | **** | **** | **** | **** | **** | **** | **** |
| Li600 | MicroNIR_NeoSpectra | **** | **** | **** | **** | **** | **** | ns |
| Li600 | NeoSpectra | **** | **** | **** | **** | **** | **** | **** |
| MicroNIR | MicroNIR_NeoSpectra | **** | *** | **** | ** | * | **** | **** |
| MicroNIR | NeoSpectra | **** | **** | *** | **** | *** | **** | **** |
| MicroNIR_NeoSpectra | NeoSpectra | **** | **** | **** | **** | * | **** | **** |

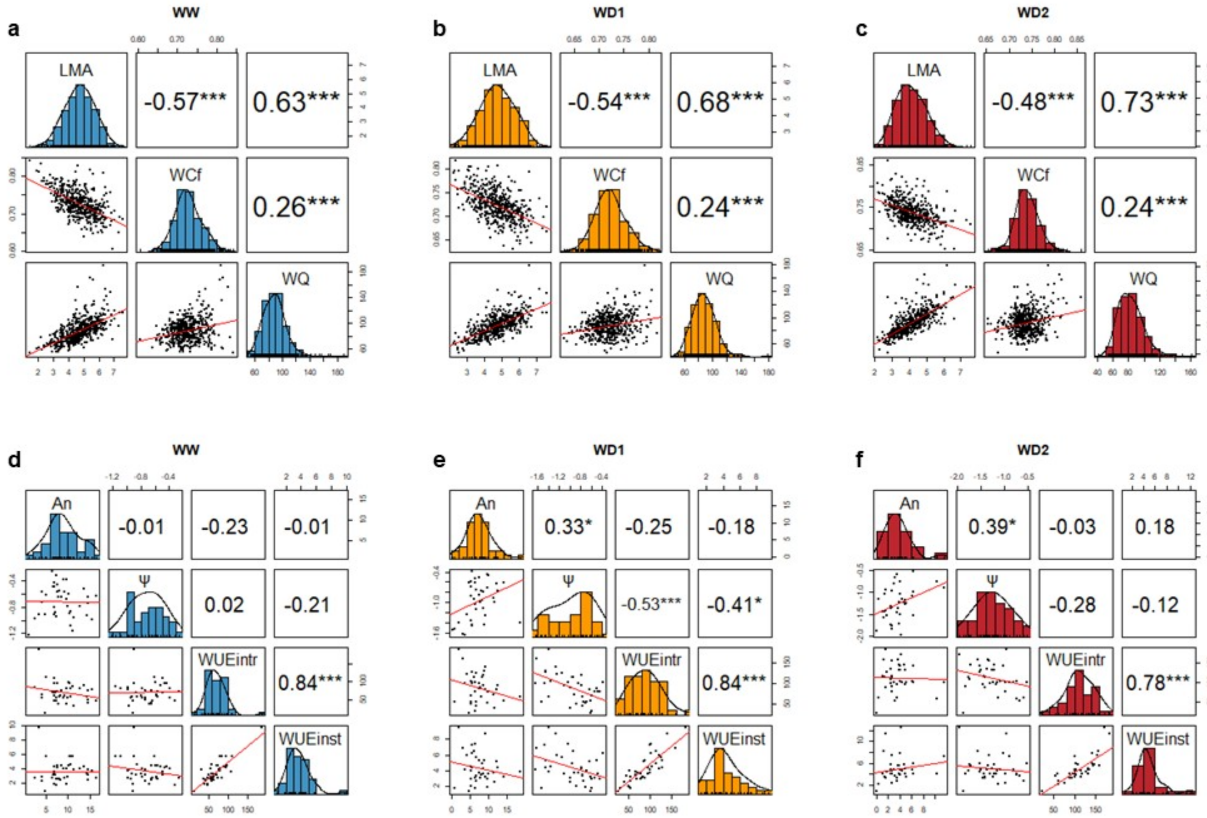


**Supplemental Figure S1. Correlation between traits directly measured with conventional methods from the greenhouse experiment, depending on the water treatment**. Correlation for leaf mass per area (LMA), water content ($WC_f$) and water quantity (WQ) measured at high-throughput for **a** Well-watered (WW), **b** Moderate water deficit (WD1) and **c** Severe water deficit (WD2) water treatment. Correlation for Net $CO_2$ assimilation ($A_n$), Leaf water potential (Ψ), Intrinsic water use efficiency ($WUE_{intr}$) and Instantaneous water use efficiency ($WUE_{inst}$) measured at low-throughput for **d** Well-watered (WW), **e** Moderate water deficit (WD1) and **f** Severe water deficit (WD2) water treatment. * = p-value ≤ 0.05; ** = p-value ≤ 0.01; *** = p-value ≤ 0.001.

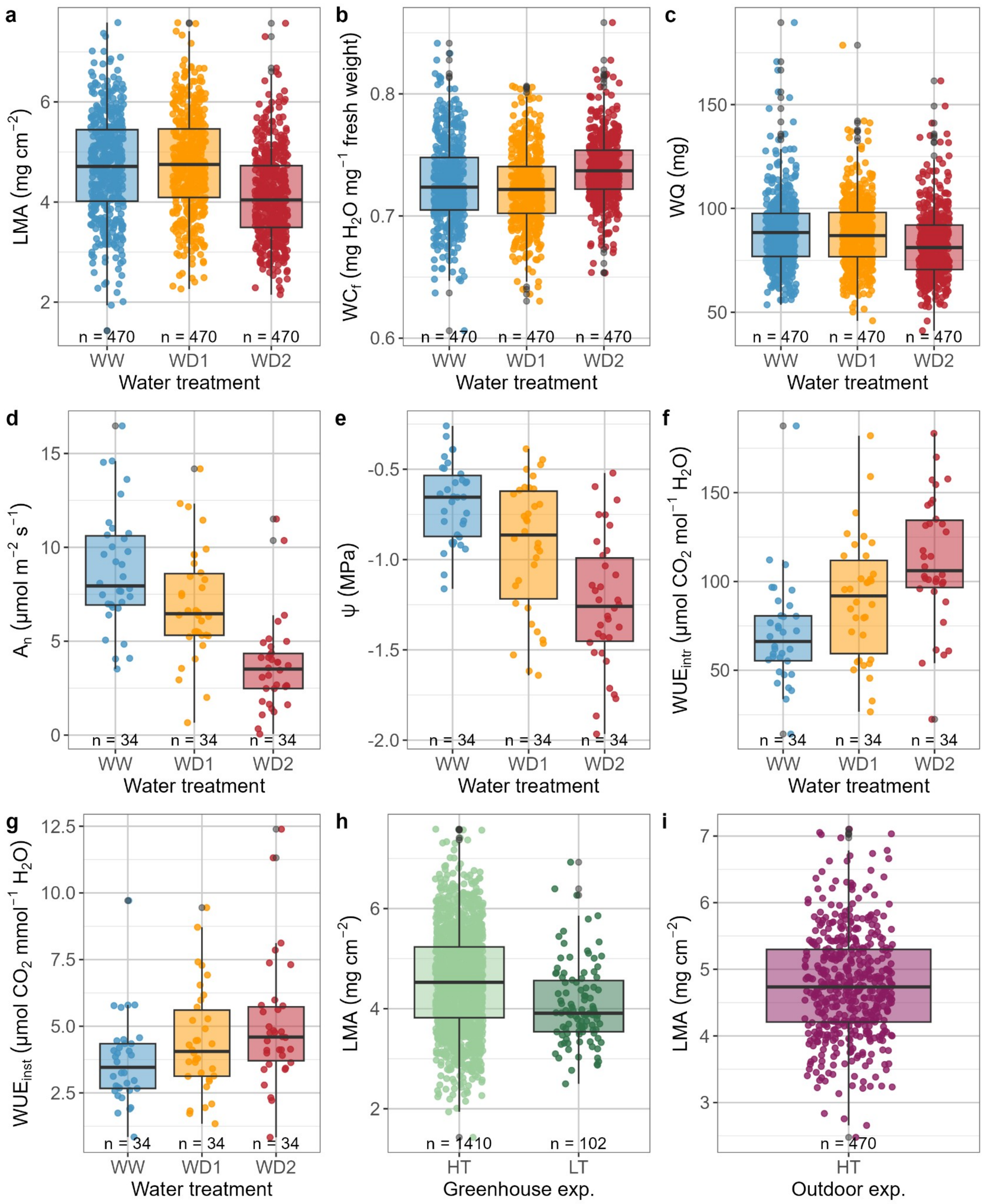


**Supplemental Figure S2. Boxplot of traits directly measured with conventional methods, depending on the water treatment or the phenotyping type. a** Leaf mass per area (LMA), **b** Water content ($WC_f$), **c** Water quantity (WQ), **d** Net $CO_2$ assimilation ($A_n$), **e** Leaf water potential

(Ψ), **f** Intrinsic water use efficiency ($WUE_{intr}$) and **g** Instantaneous water use efficiency ($WUE_{inst}$) from the greenhouse experiment with blue points and boxplot are plants in well-watered treatment (WW); orange points and boxplot are plants in moderate water deficit treatment (WD1); red points and boxplot are plants in severe water deficit treatment (WD2), **h** Leaf mass per area (LMA) according to the phenotyping type of the greenhouse experiment (HT = High-throughput, LT = Low-throughput) and **i** Leaf mass per area (LMA) from the outdoor experiment. n, number of leaves.

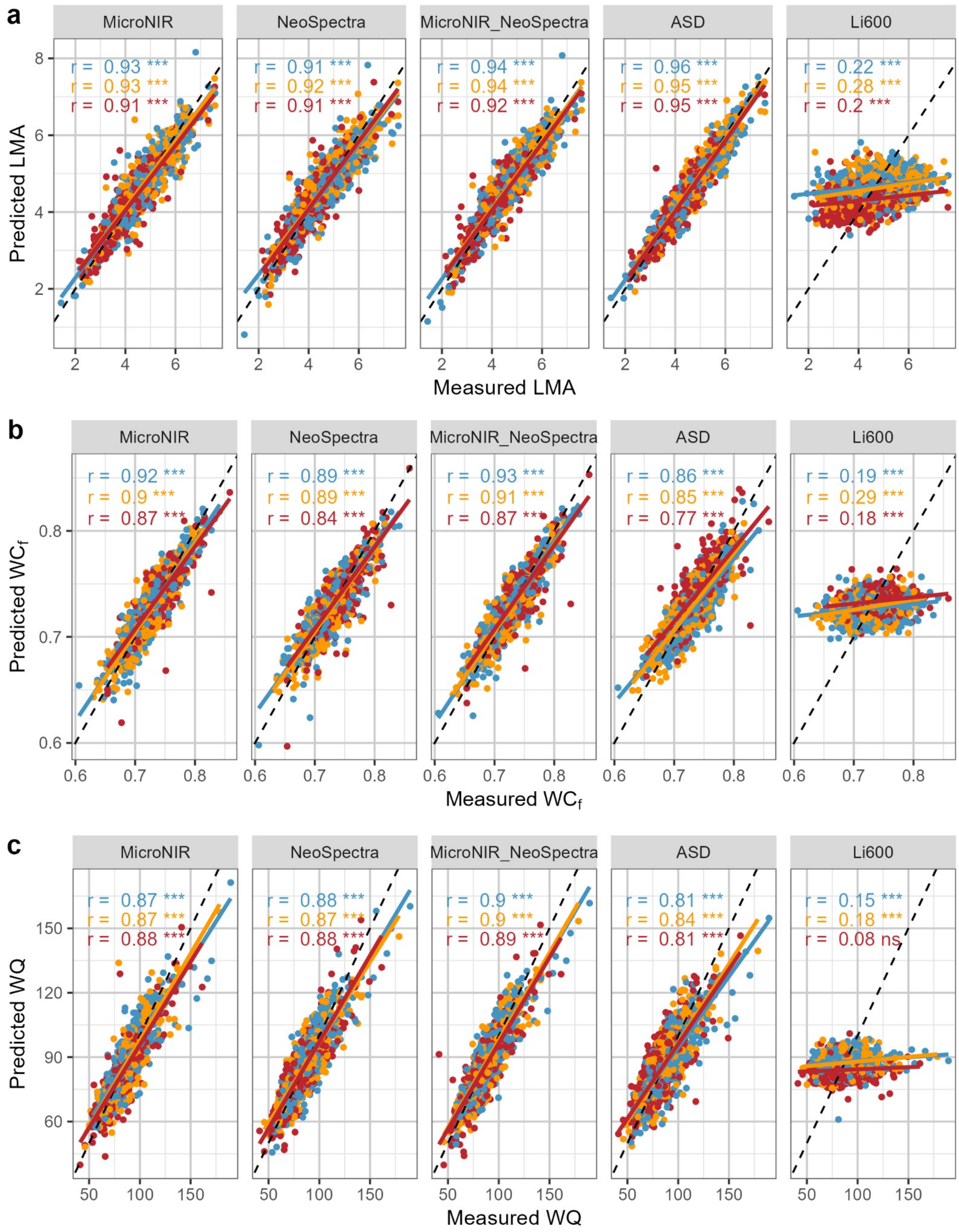
a
MicroNIR
NeoSpectra
MicroNIR_NeoSpectra
ASD
Li600
r = 0.93 ***
r = 0.93 ***
r = 0.91 ***
r = 0.91 ***
r = 0.92 ***
r = 0.91 ***
r = 0.94 ***
r = 0.94 ***
r = 0.92 ***
r = 0.96 ***
r = 0.95 ***
r = 0.95 ***
r = 0.22 ***
r = 0.28 ***
r = 0.2 ***
Predicted LMA
Measured LMA
b
r = 0.92 ***
r = 0.9 ***
r = 0.87 ***
r = 0.89 ***
r = 0.89 ***
r = 0.84 ***
r = 0.93 ***
r = 0.91 ***
r = 0.87 ***
r = 0.86 ***
r = 0.85 ***
r = 0.77 ***
r = 0.19 ***
r = 0.29 ***
r = 0.18 ***
Predicted WC$_f$
Measured WC$_f$
c
r = 0.87 ***
r = 0.87 ***
r = 0.88 ***
r = 0.88 ***
r = 0.87 ***
r = 0.88 ***
r = 0.9 ***
r = 0.9 ***
r = 0.89 ***
r = 0.81 ***
r = 0.84 ***
r = 0.81 ***
r = 0.15 ***
r = 0.18 ***
r = 0.08 ns
Predicted WQ
Measured WQ

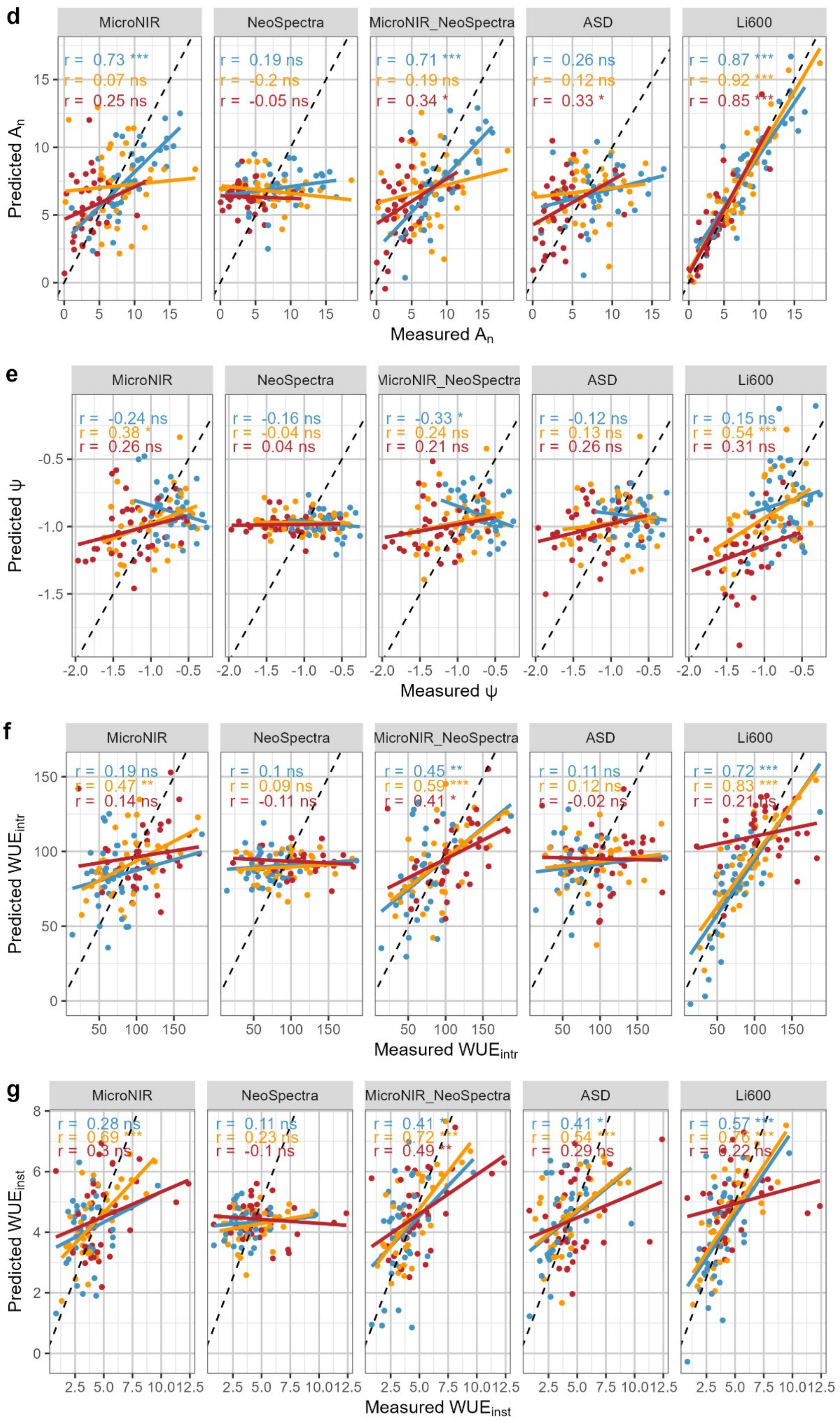
d
MicroNIR
NeoSpectra
MicroNIR_NeoSpectra
ASD
Li600
r = 0.73 ***
r = 0.07 ns
r = 0.25 ns
r = 0.19 ns
r = -0.2 ns
r = -0.05 ns
r = 0.71 ***
r = 0.19 ns
r = 0.34 *
r = 0.26 ns
r = 0.12 ns
r = 0.33 *
r = 0.87 ***
r = 0.92 ***
r = 0.85 ***
Predicted $A_n$
Measured $A_n$
e
MicroNIR
NeoSpectra
MicroNIR_NeoSpectra
ASD
Li600
r = -0.24 ns
r = 0.38 *
r = 0.26 ns
r = -0.16 ns
r = -0.04 ns
r = 0.04 ns
r = -0.33 *
r = 0.24 ns
r = 0.21 ns
r = -0.12 ns
r = 0.13 ns
r = 0.26 ns
r = 0.15 ns
r = 0.54 ***
r = 0.31 ns
Predicted ψ
Measured ψ
f
MicroNIR
NeoSpectra
MicroNIR_NeoSpectra
ASD
Li600
r = 0.19 ns
r = 0.47 **
r = 0.14 ns
r = 0.1 ns
r = 0.09 ns
r = -0.11 ns
r = 0.45 **
r = 0.59 ***
r = 0.41 *
r = 0.11 ns
r = 0.12 ns
r = -0.02 ns
r = 0.72 ***
r = 0.83 ***
r = 0.21 ns
Predicted $WUE_{intr}$
Measured $WUE_{intr}$
g
MicroNIR
NeoSpectra
MicroNIR_NeoSpectra
ASD
Li600
r = 0.28 ns
r = 0.69 ***
r = 0.3 ns
r = 0.11 ns
r = 0.23 ns
r = -0.1 ns
r = 0.41 *
r = 0.72 ***
r = 0.49 **
r = 0.41 *
r = 0.54 ***
r = 0.29 ns
r = 0.57 ***
r = 0.76 ***
r = 0.22 ns
Predicted $WUE_{inst}$
Measured $WUE_{inst}$

**Supplemental Figure S3. Cross-validation observed versus predicted values for each trait predicted by each device (datasets A and B in Table 1). a** Leaf mass per area (LMA), **b** Water content ($WC_f$), **c** Water quantity (WQ), **d** Net $CO_2$ assimilation ($A_n$), **e** Leaf water potential (Ψ), **f** Intrinsic water use efficiency ($WUE_{intr}$) and **g** Instantaneous water use efficiency ($WUE_{inst}$). The blue dots are plants in well-watered treatment (WW); orange dots are plants in moderate water deficit treatment (WD1); red dots are plants in severe water deficit treatment (WD2). The black dashed line is 1:1 line. r is the correlation coefficient. ns = p-value ≥ 0.05; * = p-value ≤ 0.05; ** = p-value ≤ 0.01; *** = p-value ≤ 0.001.

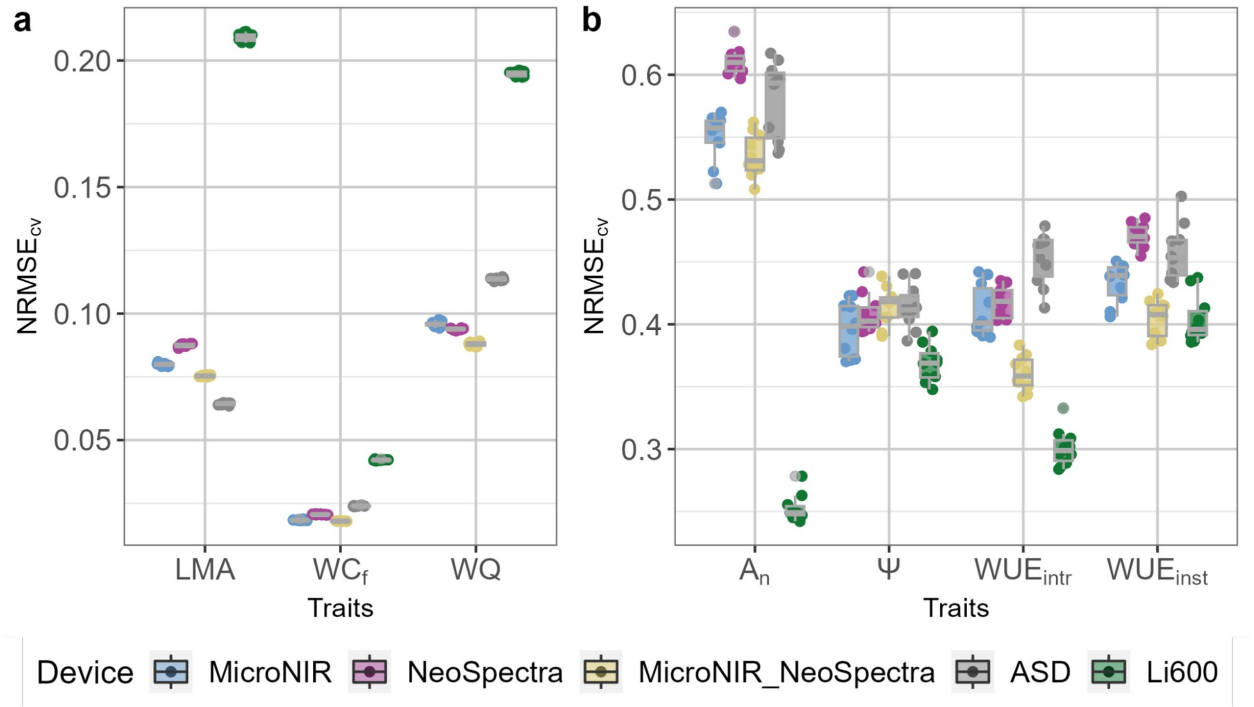


**Supplemental Figure S4. Cross-validation normalized root mean square error (NRMSE) of predicted traits according to the device. a** High-throughput measurements of Leaf mass per area (LMA), Water content ($WC_f$) and Water quantity (WQ) (dataset A in Table 1). **b** Low-throughput measurements of Net $CO_2$ assimilation ($A_n$), Leaf water potential (Ψ), Intrinsic water use efficiency ($WUE_{intr}$) and Instantaneous water use efficiency ($WUE_{inst}$) (dataset B in Table 1).

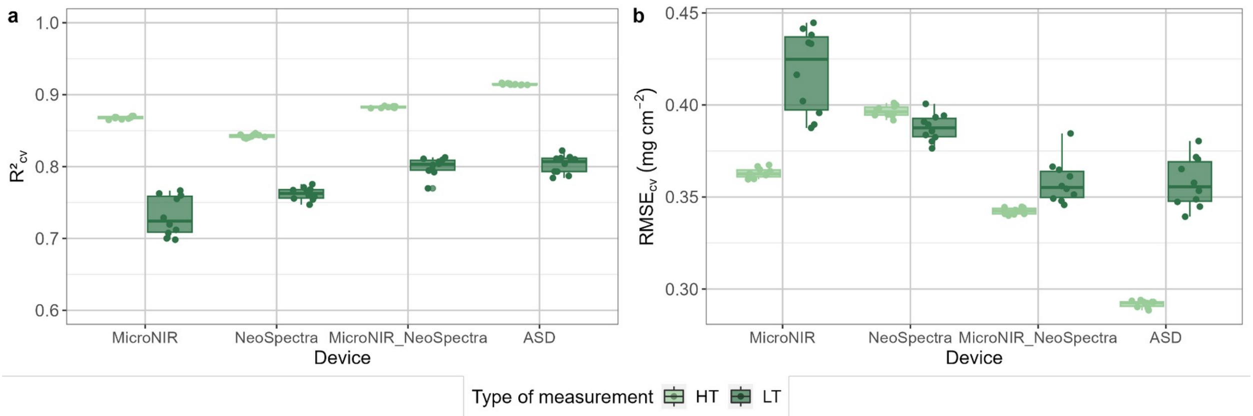


**Supplemental Figure S5. Cross-validated prediction quality of leaf mass per area (LMA) prediction using low- (109 leaves, dataset B in Table 1) or high-throughput (1437 leaves, dataset A in Table 1) measurements in the calibration set, according to the device.** (A) Corresponds to the $R^2_{cv}$ and (B) corresponds to the $RMSE_{cv}$ expressed in mg $cm^{-2}$. Each point in the boxplot corresponds to one of the 10 iterations of the cross-validation. For each device, the low- and high-throughput $R^2_{cv}$ and $RMSE_{cv}$ are significatively different (p-value < 0.01).

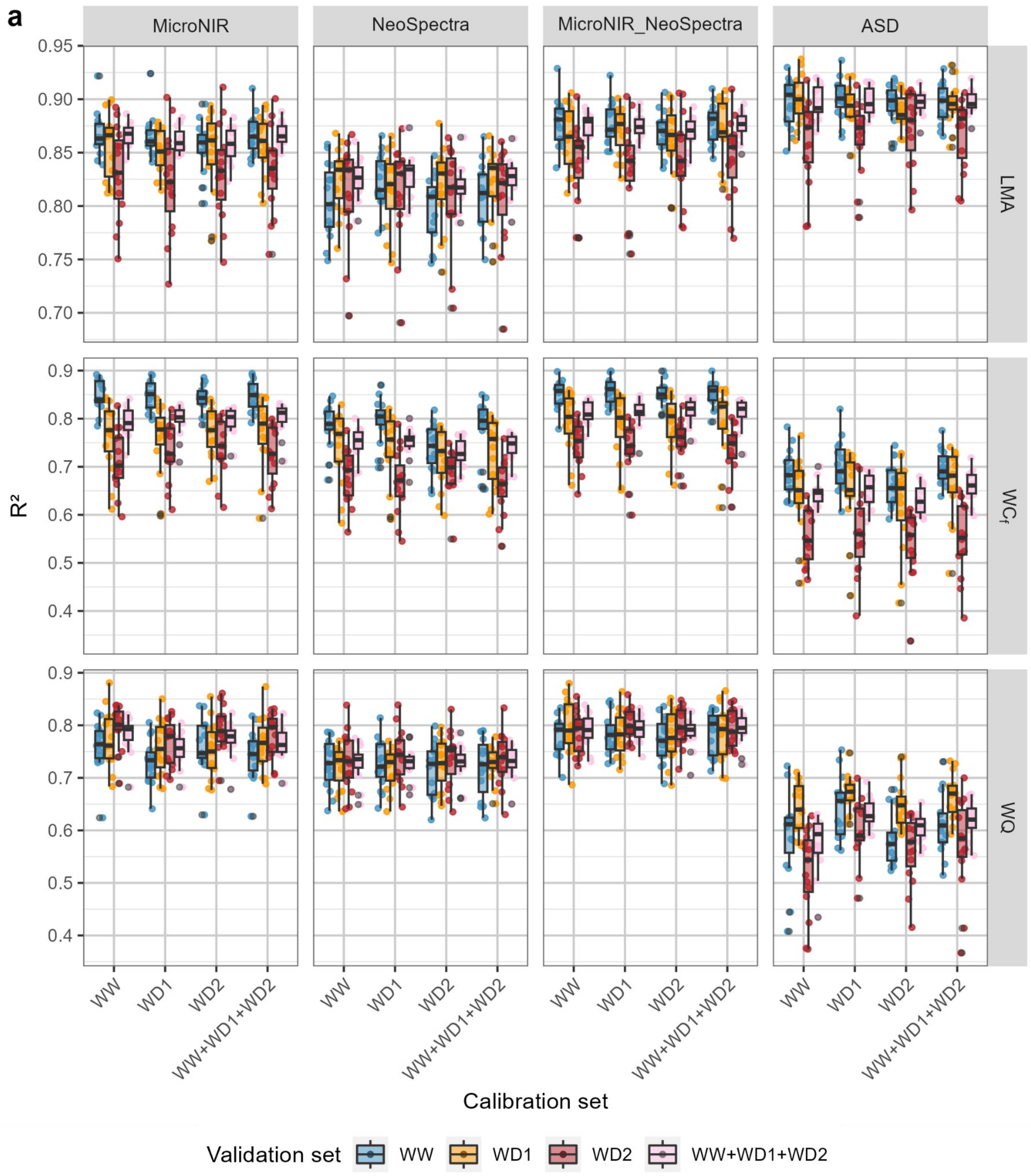
a
MicroNIR
NeoSpectra
MicroNIR_NeoSpectra
ASD
LMA
WC$_f$
WQ
R²
Calibration set
WW
WD1
WD2
WW+WD1+WD2
Validation set
WW
WD1
WD2
WW+WD1+WD2

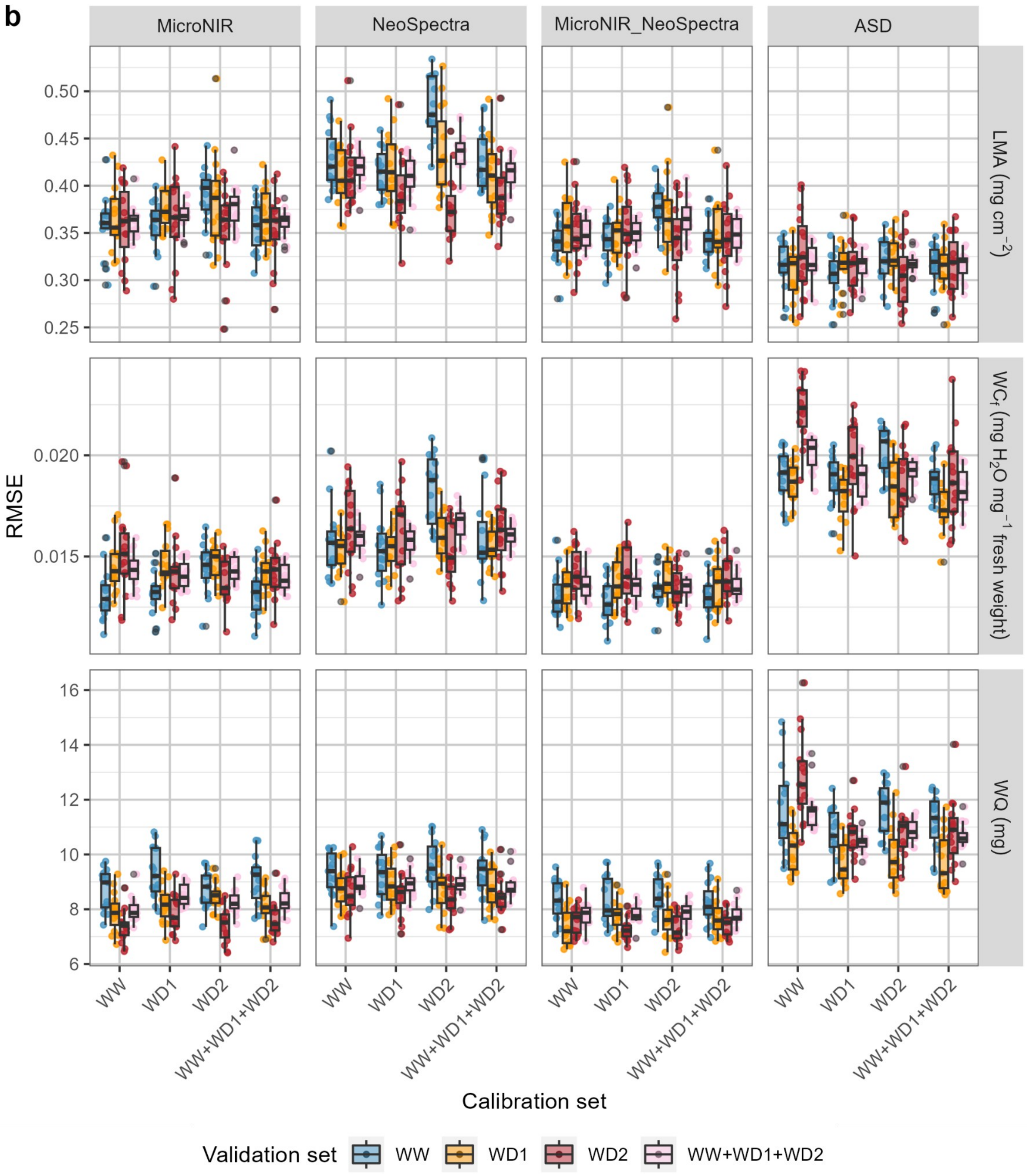


**Supplemental Figure S6. Prediction quality of PLSR models for leaf mass per area (LMA), water content ($WC_f$) and water quantity (WQ) depending on the water treatment validation sets predicted by the WW calibration set (dataset F in Table 1), WD1 calibration set (dataset G in Table 1), WD2 calibration set (dataset H in Table 1) and WW+WD1+WD2 calibration set**

**(dataset I in Table 1) according to the device. a** corresponds to the $R^2$ and **b** corresponds to the RMSE of the prediction. Each point in the boxplot corresponds to one of the 15 iterations of the validation (as detailed in Table 1). The blue dots are plants in well-watered treatment (WW); orange dots are plants in moderate water deficit treatment (WD1); red dots are plants in severe water deficit treatment (WD2).

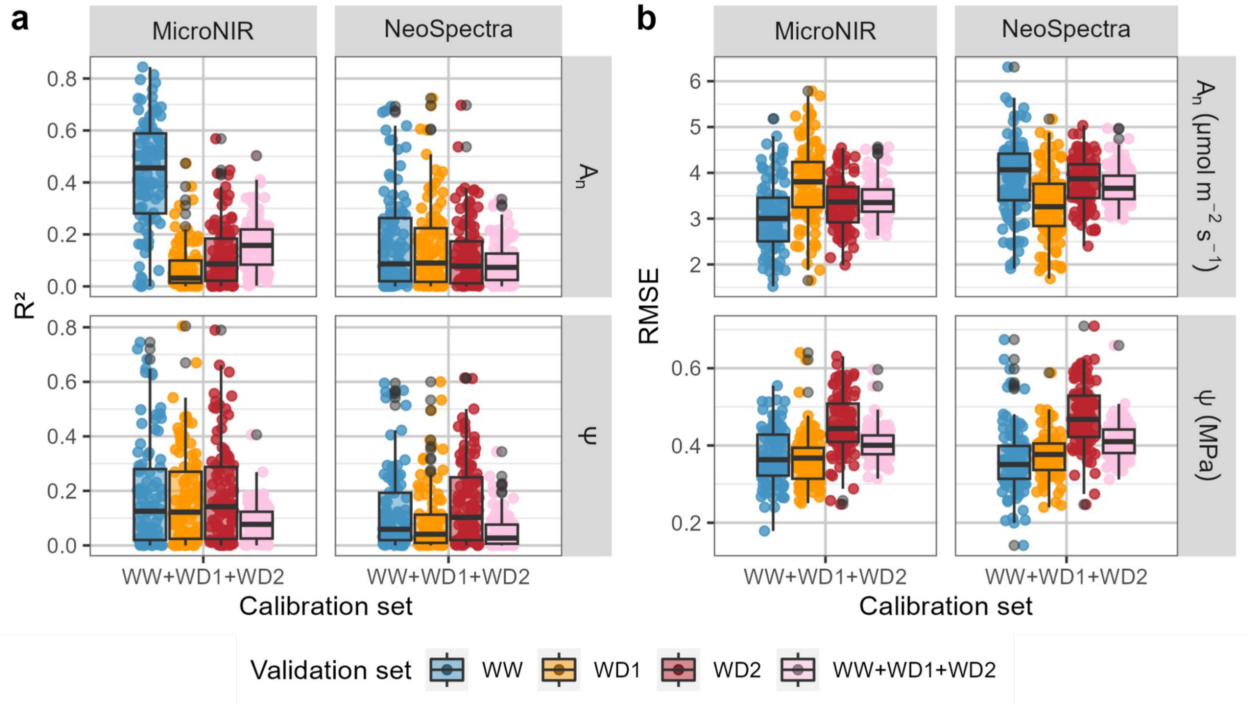


**Supplemental Figure S7. Prediction quality of PLSR models for Net $CO_2$ assimilation ($A_n$) and Leaf water potential (Ψ) depending on the water treatment validation sets and according to the device.** Models are calibrated using the WW+WD1+WD2 calibration set (dataset J in Table 1). **a** corresponds to the $R^2$ and **b** corresponds to the RMSE of the prediction. Each point in the boxplot corresponds to one of the 100 iterations of the validation (as detailed in Table 1). The blue dots are plants in WW treatment; orange dots are plants in WD1 treatment; red dots are plants in WD2 treatment; pink dots are all WW, WD1 and WD2 plants.

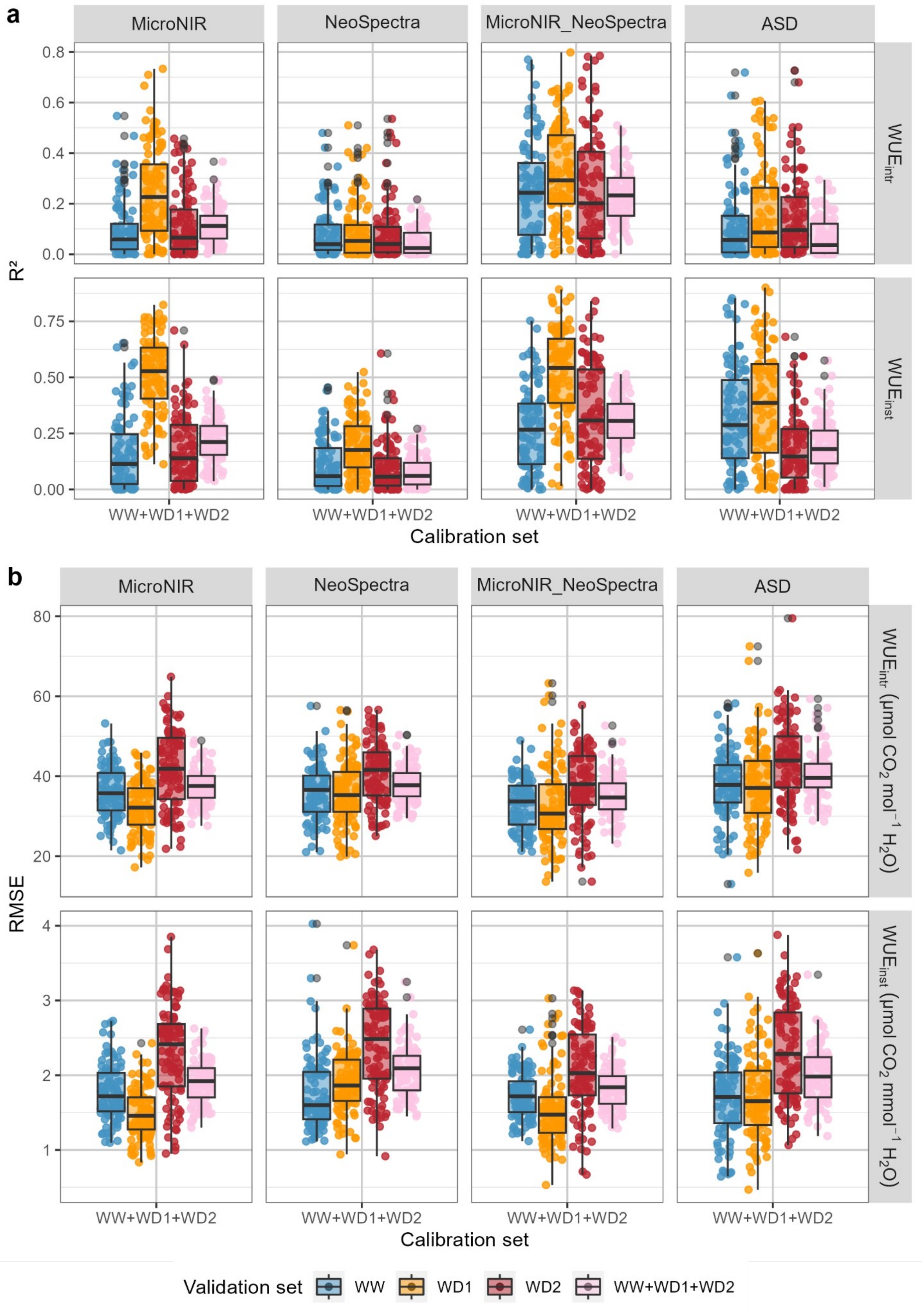
a
MicroNIR
NeoSpectra
MicroNIR_NeoSpectra
ASD
R²
$WUE_{intr}$
$WUE_{inst}$
WW+WD1+WD2
Calibration set
b
RMSE
$WUE_{intr}$ ($\mu mol\ CO_2\ mol^{-1}\ H_2O$)
$WUE_{inst}$ ($\mu mol\ CO_2\ mmol^{-1}\ H_2O$)
Validation set
WW
WD1
WD2
WW+WD1+WD2

**Supplemental Figure S8. Prediction quality of Intrinsic water use efficiency ($WUE_{intr}$) and Instantaneous water use efficiency ($WUE_{inst}$) prediction according to the water treatment validation set predicted by the WW+WD1+WD2 calibration set (dataset J in Table 1) according to spectrometer devices. a** corresponds to the $R^2$ and **b** corresponds to the RMSE of the prediction. Each point in the boxplot corresponds to one of the 100 iterations of the validation (as detailed in Table 1). The blue dots are plants in WW treatment; orange dots are plants in WD1 treatment; red dots are plants in WD2 treatment; pink dots are all WW, WD1 and WD2 plants.

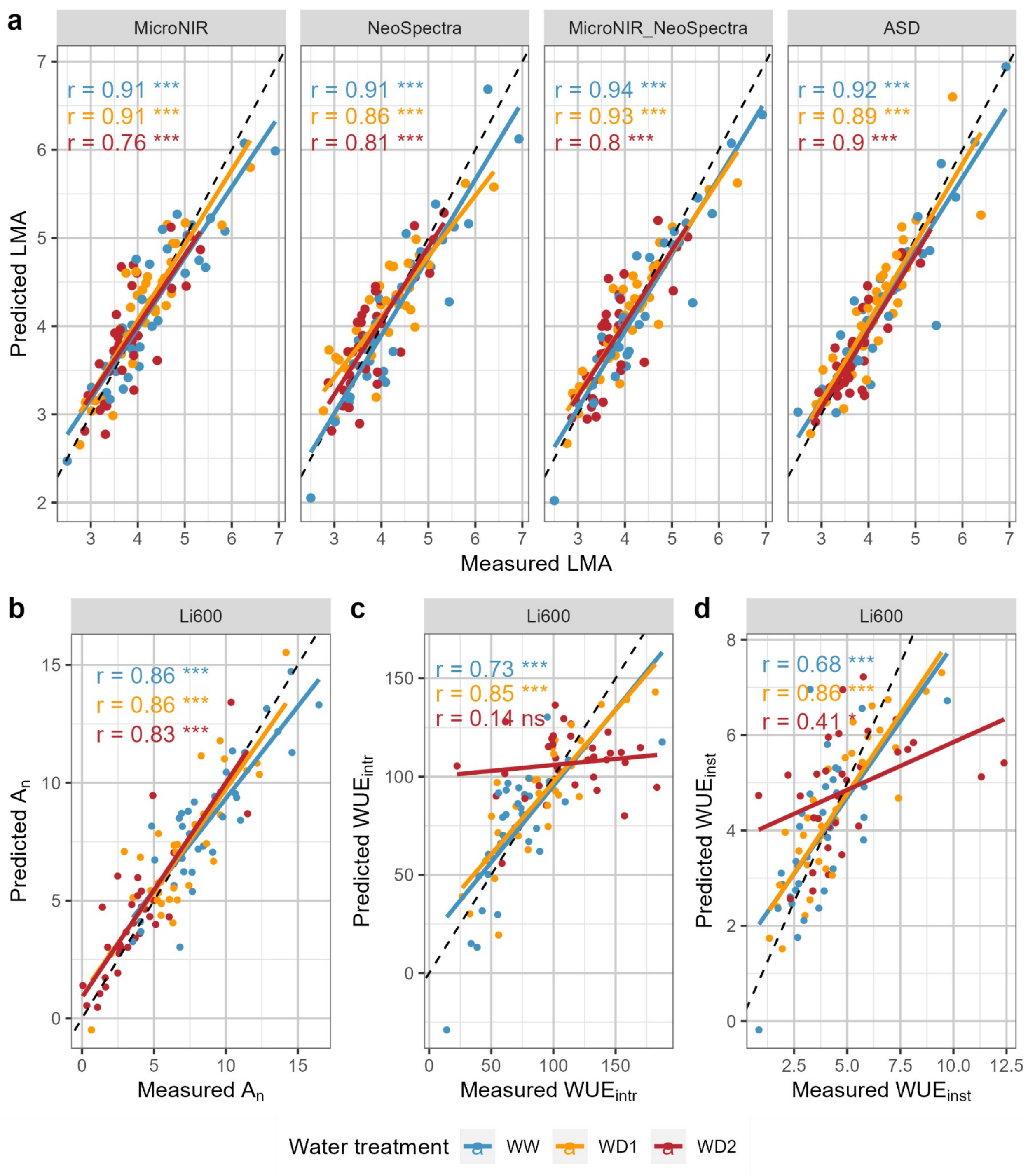


**Supplemental Figure S9. Cross-validation observed *versus* predicted values (dataset K in Table 1)** of **a** Leaf mass per area (LMA) predicted with spectrometers (MicroNIR, NeoSpectra, MicroNIR_NeoSpectra and ASD), **b** Net $CO_2$ assimilation ($A_n$), **c** Intrinsic water use efficiency ($WUE_{intr}$) and **d** Instantaneous water use efficiency ($WUE_{inst}$) predicted with Li600. The blue dots are plants in well-watered treatment (WW); orange dots are plants in moderate water deficit treatment (WD1); red dots are plants in severe water deficit treatment (WD2). r is the correlation coefficient. ns = p-value ≥ 0.05; * = p-value ≤ 0.05; ** = p-value ≤ 0.01; *** = p-value ≤ 0.001.
